# ATP13A1 engages GET3 to facilitate substrate-specific translocation

**DOI:** 10.1101/2024.02.12.579870

**Authors:** Xiaoyan Yang, Tingting Li, Zhiyu Fang, Zhigang Feng, Yan Zou

## Abstract

Proper localization of proteins to their final destinations is crucial for preserving cellular structure and functions. The interpretation and sorting of highly variable targeting sequences in secreted and membrane proteins, however, pose a challenge in achieving precise localization within specific secretory apparatus. In this study, we demonstrate that atypical signal sequences characterized by high hydrophobicity and/or the absence of characteristic charges undergo targeting to the endoplasmic reticulum (ER) in a reverse orientation, followed by partial cleavage. The P5A- ATPase ATP13A1 recognizes the cleaved signal sequence and dislocates it to the targeting factor GET3, subsequently engaging SEC61 for further translocation. Our findings unveil a comprehensive translocation pathway that operates in a substrate- specific manner, ensuring both high efficiency and fidelity in the protein subcellular localization.

## Introduction

Accurate protein localization to their respective destinations is crucial for maintaining cellular structure and homeostasis. Approximately one-third of the eukaryotic proteome comprises secreted and membrane proteins designated for the intercellular space or intracellular organelles (Uhlen et al., 2015). Throughout biogenesis, a majority of these proteins undergo co- or post-translational translocation into the endoplasmic reticulum (ER) to traverse its impermeable membrane (Ast et al., 2013; Ng et al., 1996; Rapoport et al., 2017).

The process of ER translocation is intricate and tightly regulated, involving two major machineries. Initially, the N-terminal signal sequences (SSs) or transmembrane helices (TMs) of translating polypeptides are recognized by the signal recognition particle (SRP) in the cytosol and subsequently delivered to the SRP receptor (SRPR) for ER targeting (Akopian et al., 2013). The nascent polypeptides are then loaded onto the SEC61 translocon and guided into the ER lumen until stop signals, such as transmembrane helices, emerge from ribosomes. Following import, SSs are promptly removed from the translocated polypeptides by the signal peptide peptidase (SPP) (Blobel and Dobberstein, 1975; Weihofen et al., 2002). In contrast to the SRP complex, post-translationally translocated proteins utilize chaperones to prevent their folding in the cytosol, along with Sec62-Sec63 to facilitate their translocation (Rapoport et al., 2017).

Defects in the translocation machineries have been associated with various developmental deficits and diseases. For instance, mutations in the SRP complex lead to syndromic neutropenia (Carapito et al., 2017) and familial aplasia and myelodysplasia (Kirwan et al., 2012). Dysregulation of factors in the SEC61 complex is linked to kidney and liver diseases, diabetes, and human cancer (Linxweiler et al., 2017). Apart from the SEC61 translocon, the ER membrane protein complex (EMC) and the GET complex (Guna et al., 2018; Mariappan et al., 2011; Stefanovic and Hegde, 2007) also play roles in the ER targeting of specific SSs and tail-anchored (TA) proteins. Targeting sequences exhibit high variability and degeneracy, evolving to specialize in targeting different organelles through distinct targeting or translocation machinery (Owji et al., 2018). Nevertheless, the precise mechanisms by which distinct targeting sequences are interpreted and sorted by the secretory apparatus in a high-fidelity manner remain elusive.

P-type ATPases forma phylogenetically conserved superfamily responsible for transporting various substrates, including ions, phospholipids, and polyamines, across membranes (Huang et al., 2022; Palmgren and Nissen, 2011). Among the P-type ATPase family, the P5A ATPase, referred to as ATP13A1, stands out as a recently identified dislocase to remove mislocalized TA proteins from the ER, preventing their degradation through ER-associated degradation (ERAD) pathways (McKenna et al., 2022; McKenna et al., 2020; Qin et al., 2020). Moreover, ATP13A1 has been implicated in the SS-mediated ER translocation of secreted and membrane proteins (Feng et al., 2020; Li et al., 2021). Quantitative proteomics studies support the essential role of ATP13A1 in maintaining the stability of N-terminal ER-targeting SS/TM proteins and TA proteins, with N-terminal SS proteins being the largest class destabilized by *ATP13A1* knockout (KO) (McKenna et al., 2020).

Physiological investigations demonstrate that P5A ATPases exert pleiotropic effects in metazoans through various essential substrates. In *C. elegans*, null mutations in *catp-8*, the ortholog of ATP13A1, impact the ER translocation of neuronal receptor DMA-1 and morphogen Wnt proteins, leading to effects on intestinal development and lifespan (Feng et al., 2020; Li et al., 2021; Qin et al., 2020; Tang et al., 2021). In mice, ATP13A1 is crucial for the stability of antiviral proteins MAVS. Atp13a1 knockout mice exhibit growth retardation and embryonic lethality, indicating the significance of P5A ATPases in diverse physiological processes (Zhang et al., 2022).

Furthermore, a hypomorphic mutation in human ATP13A1 is associated with a range of developmental defects, including intellectual disability, attention deficit hyperactivity disorder, and facial and nail abnormalities (Anazi et al., 2017; Tang et al., 2021). However, existing studies primarily focus on the fate and pathological consequences of N-terminal SS-containing proteins in *ATP13A1* KO cells, leaving unanswered questions regarding how ATP13A1 mediates the ER translocation of SS-containing proteins under physiological conditions.

In this study, we employed a combination of bulk fluorescence, biochemical, and genetic assays to investigate ATP13A1’s role in SS-mediated ER translocation. Our findings reveal that the topogenesis of atypical SSs, characterized by unusually high hydrophobic cores and/or lacking N-terminal positively charged residues, is error- prone. These atypical SSs undergo cleavage when targeted to the ER membrane through the SRP complex. Subsequently, the partially cleaved SSs are dislocated by ATP13A1 from the ER membrane to the targeting factor GET3 and then conducted into the ER through the SEC61 complex. Our study elucidates a comprehensive translocation pathway that operates in a substrate-specific manner to maintain high efficiency and fidelity. The emergence of P5A ATPases in eukaryotes addresses the evolving demands of signal sequences, which diverge evolutionarily to target different organelles.

## Results

### ER Translocation of SSs with High Hydrophobicity and/or Lacking N-terminal Positively Charged Residues Mediated by P5A ATPase

To investigate the regulation of ER translocation *in vivo* by P5A ATPase, we first developed a DMA-1SS (Feng et al., 2020) reporter system, comprising DMA- 1SS::moxDendra2::Opsin::FLAG, to assess its translocation efficiency into the ER. mCherry was co-expressed as an internal control using SL2 trans-splicing within the same operon (Figure 1A). As anticipated, the DMA-1SS reporter, but not mCherry, was secreted into the pseudocoelom and predominantly taken up by coelomocytes in wild-type (WT) animals. In contrast, the DMA-1SS reporter was retained in intestinal cells in *catp-8/P5A* mutants (Figure 1B). Utilizing DMA-1ΔTM as a positive control, given its failure to be translocated into the ER and degradation in *catp-8 mutants* (Feng et al., 2020), we validated that the DMA-1SS reporter mimicked the behavior of DMA-1ΔTM, failing to be N-glycosylated in *catp-8* mutants (Figures 1C and 1D). Therefore, the DMA-1SS reporter serves as a sensitive indicator of the ER translocation of P5A-dependent SSs.

**Figure 1.**
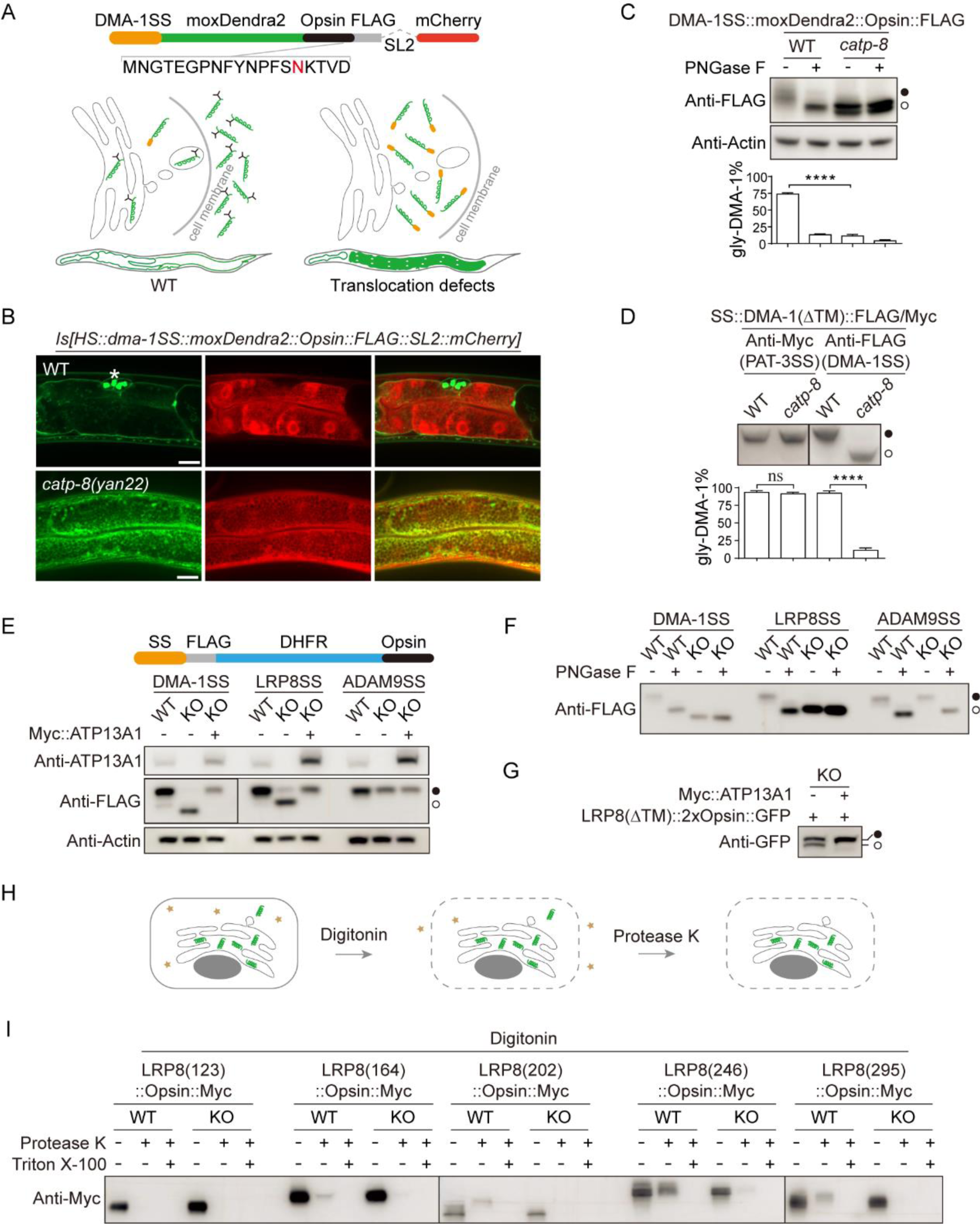
P5A is required for SS-mediated ER translocation. |(A) Diagram of DMA-1SS::moxDendra2::Opsin::FLAG::SL2::mCherry reporter (ZOU908 and ZOU909 Strains) to assay P5A-dependent translocation efficiency. The Opsin tag in this reporter is N-glycosylated when it is secreted into the pseudocoelom from the intestine of WT animals. It is unglycosylated when it is trapped inside the intestinal cells of translocation-deficient mutants. (B) Secretion of DMA-1SS::moxDendra2::Opsin::FLAG is disrupted in *catp-8* mutants. Representative images showing the moxDendra2 localization of DMA- 1SS::moxDendra2::Opsin::FLAG in WT and *catp-8* worms. Asterisk indicates coelomocytes. Scale bars, 10 μm. (C) N-glycosylation of DMA-1SS::moxDendra2::Opsin::FLAG is abolished in *catp-8* mutants. Extracted proteins were incubated with the PNGase F enzyme to remove N- linked oligosaccharides in the Opsin tag. When fully translocated into the ER, the glycosylated opsin tag generates a slowly-migrating form as indicated by the black circle. In contrast, translocation deficient proteins shows an unglycosylated and fast- migrating band as indicated by the clear circle. (D) CATP-8 is specifically required for ER translocation of DMA-1SS but not PAT- 3SS. DMA-1SS::DMA-1ΔTM::FLAG and PAT-3SS::DMA-1ΔTM::Myc were integrated into WT (ZOU977 strain) and then crossed into *catp-8* worms (ZOU979 strain). (E) ATP13A1 is specifically required for ER translocation of LRP8SS but not ADAM9SS. SS::FLAG::DHFR::Opsin reporter was used to assay the translocation efficiency of DMA-1SS, LRP8SS, or ADAM9SS in WT, *ATP13A1* KO, or KO cells rescued with ATP13A1. (F) Incubation with the PNGase F enzyme to remove the N-linked oligosaccharides validates that the differences in band sizes for (E) were due to N-glycosylation. (G) ATP13A1 is required for ER translocation of LRP8(ΔTM) proteins. (H) Diagram illustrating biochemical protease protection (BPP) experiment. In the BPP assay, the cholesterol-rich plasma membrane is selectively penetrated by digitonin, a cholesterol-binding drug, to allow protease K to enter the cytosol. Therefore, cytosolic proteins in the semi-permeabilized cells are digested by protease K, while proteins within low cholesterol organelles, such as the ER, are still present. (I) LRP8(123) is protease accessible in the BPP assay. LRP8 truncation at different lengths (123, 164, 202, 246, and 295 amino acids) were transfected into WT or *ATP13A1* KO cells, followed by BPP treatment and western blot. Quantifications of glycosylated proteins are from 3 biological replicates and presented as means ± SEMs; ns, not significant (*p* > 0.05); \*\*\*\**p* < 0.0001 (Student’s *t*-test). Black and clear circles in (C)-(G) indicate the N-glycosylated and the unglycosylated proteins, respectively. See also Figure S1.

In addition to DMA-1SS from *C. elegans*, we screened various SSs whose protein abundance is contingent on ATP13A1 in mass spectrometry (MS) analysis (McKenna et al., 2020). Our findings revealed that the ER translocation of multiple SSs, including LRP8SS, necessitated the presence of the P5A ATPase (Figures 1E-1G, and S1A). Consequently, we inferred that P5A-dependent SSs exhibit high hydrophobicity and/or lack N-terminal positively charged residues (Figure S1B). For subsequent investigations into P5A-mediated translocation, we selected DMA-1SS from *C. elegans* and human LRP8SS as positive representatives. As negative controls, we chose PAT-3SS from *C. elegans* (Feng et al., 2020) and human ADAM9 SS, both of whose translocation is independent of P5A ATPase (Figures 1E, 1F, and S1A).

Remarkably, in comparison to the full length and long truncations of LRP8SS, we observed that LRP8SS with a short passenger protein of 123 amino acids, LRP8 (123), failed to enter the ER (Figures 1G-1I), suggesting that the ER translocation of LRP8SS may necessitate molecular chaperones recruited by passenger proteins.

### ATP13A1 Interacts Directly with SSs through its Substrate Binding Pocket

Our hypothesis posits that the P5A-dependent SSs represent a subclass of substrates for ATP13A1. However, capturing their interactions through co-immunoprecipitation (Co-IP) proves challenging due to the transient nature of the process. As an alternative, we employed the PUP-IT proximity tagging system (Liu et al., 2018) and observed physical proximity betweenLRP8SS and PafA::ATP13A1 (Figures S2A and S2B). To validate this finding, we utilized a superfolder Green Fluorescence Protein (sfGFP) fused with LRP8SS to impede the translocation process, resulting in a translocation intermediate (Ma et al., 2019). As anticipated, the bulky sfGFP diminished the translocation efficiency of LRP8SS into the ER, evidenced by reduced glycosylation in WT cells (Figure S2C). Further investigation of the subcellular localization of LRP8SS::sfGFP, achieved through co-transfection with a GFPnano::RFP reporter in HEK293FT cells, revealed a change in localization, validating the slowed LRP8SS translocation (Figure 2A). Selective penetration of the cholesterol-rich plasma membrane with digitonin and subsequent removal of cytosolic proteins demonstrated the enrichment of the GFPnano::RFP reporter on the ER membrane, confirming sfGFP’s effect on LRP8SS translocation (Figure 2A). With the assistance of sfGFP, we identified that endogenous ATP13A1 specifically co- immunoprecipitated with LRP8SS but not the P5A-independent PAT-3SS (Figure 2B). To delve deeper into the interaction sites between ATP13A1 and LRP8SS, we induced an ATP-bound state by utilizing the D533N hydrolysis mutant of ATP13A1 to stabilize its interaction with LRP8SS. Additionally, we employed an orthogonal system using the suppressor tRNA/aminoacyl-tRNA synthetase (aaRS) pair for the photolabile non-canonical amino acid (ncAA) p-azido-L-phenylalanine (pAzF) (Figures 2C and S2D) to crosslink residues between interacting proteins at a distance range of 3–4 Å (Gautier et al., 2010). Mapping the insertion position of pAzF in LRP8SS through an amber stop codon (TAG), we determined that LRP8(25TAG)SS, while affecting neither the expression level nor the P5A-dependence of LRP8SS (Figure S2E), could be used to crosslink its interacting proteins. Upon ultraviolet (UV) illumination, LRP8(25TAG)SS crosslinked with ATP13A1(D533N) (Figure 2D). Cryo–electron microscopy structures unveiled the putative substrate binding pocket of Spf1, the P5A ortholog in *Saccharomyces cerevisiae* (McKenna et al., 2020). Consequently, we selected corresponding residues (Y265, M272, F276, E492, I495, L499, A1025) in the putative substrate binding pocket of ATP13A1 to incorporate pAzF. Among these residues, ATP13A1 crosslinked with LRP8SS at M272 and L499 (Figures 2E and 2F). Thus, these findings suggest that ATP13A1 establishes a direct interaction with LRP8SS through its substrate binding pocket.

**Figure 2.**
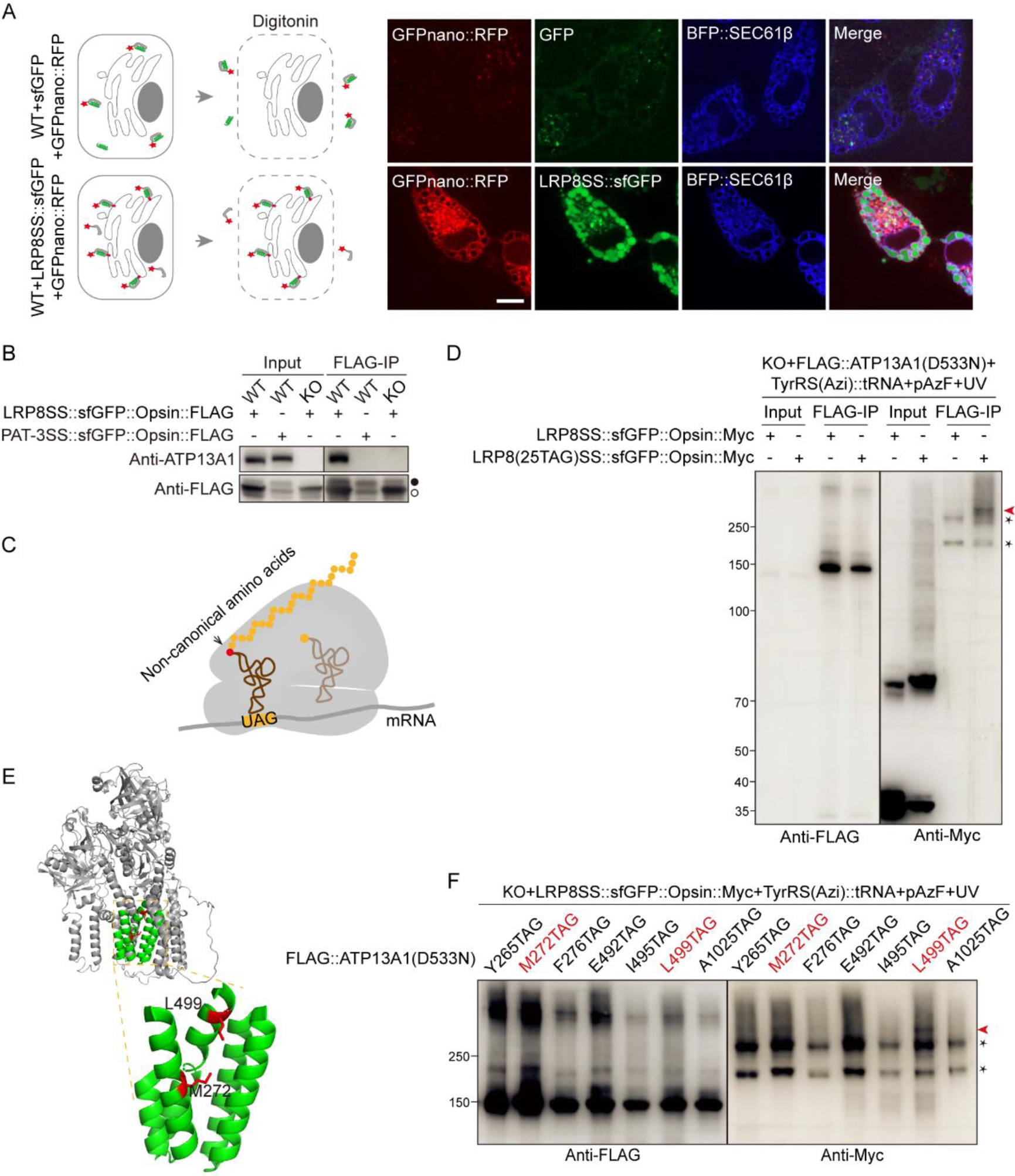
P5A ATPase directly interacts with SS through its substrate binding pocket. (A) Diagram and representative images of a GFPnano::RFP reporter to assay the subcellular localization and topology of LRP8SS. RFP was fused with the GFP nanobody (GFPnano) and co-expressed with sfGFP or LRP8SS::sfGFP in HEK293FT cells. Cytosolic sfGFP and GFPnano::RFP were released outside through digitonin penetrated cell membrane in the assay. In contrast, LRP8SS::sfGFP was inserted into the ER membrane with sfGFP facing cytosol, which bound cytosolic GFPnano::RFP to prevent it from flowing out of the digitonin penetrated cell membrane. ER was labeled with TagBFP::SEC61β. Scale bar, 5 μm. (B) Endogenous ATP13A1 specifically co-immunoprecipitated with LRP8SS but not PAT-3SS. LRP8SS::sfGFP::Opsin::FLAG or PAT-3SS::sfGFP::Opsin::FLAG was expressed in WT and *ATP13A1* KO cells. Black and clear circles indicate the N- glycosylated and unglycosylated proteins, respectively. (C) Diagram of the orthogonal system which uses the suppressor tRNA/aminoacyl- tRNA synthetase to incorporate photolabile ncAA with an amber stop codon at targeting positions. (D) LRP8SS is crosslinked with ATP13A1. LRP8(25TAG)SS::sfGFP::Opsin::Myc or LRP8SS::sfGFP::Opsin::Myc was co-transfected with FLAG::ATP13A1(D533N) in *ATP13A1* KO cells. (E) Illustration of tested sites (red residues) in the substrate binding pocket (green helixes) of ATP13A1. The ATP13A1 structure was inferred from the yeast ortholog of P5A ATPase, SPF1(McKenna et al., 2020). (F) Photo-crosslinking analysis surveying the interacting residues of ATP13A1 with LRP8SS. The red arrow represents the ATP13A1 band crosslinked with LRP8SS. Asterisks indicate non-characteristic bands caused by endogenous proteins with TAG stop codon. See also Figure S2.

### ATP13A1 Facilitates ER Translocation of P5A-dependent SSs with Incorrect, Inverted Topology

While depletion of *catp-8* resulted in the failure of ER translocation for DMA- 1ΔTM (Figure 1D), DMA-1ΔTM::GFP was observed surrounding the ER marker, rather than dispersing in the cytosol in *catp-8* mutant worms (Figure 3A). Correspondingly, knocking out *ATP13A1* led to the ER localization of GFPnano::RFP, indicating that LRP8SS::GFP, with its C-terminal GFP in the cytosol, attached to the ER in *ATP13A1* KO cells (Figure 3B). In contrast, LRP8SS::GFP entered the ER, causing GFPnano::RFP to be evenly distributed in the cytosol in WT cells or KO cells rescued with ATP13A1 (Figure 3B). Furthermore, in *ATP13A1* KO cells, LRP8SS was membrane-integrated, as evidenced by its resistance to extraction with sodium carbonate (Figure 3C). Taken together, these results imply that *ATP13A1* depletion leads to the ER targeting of P5A-dependent SSs in an incorrect topology.

**Figure 3.**
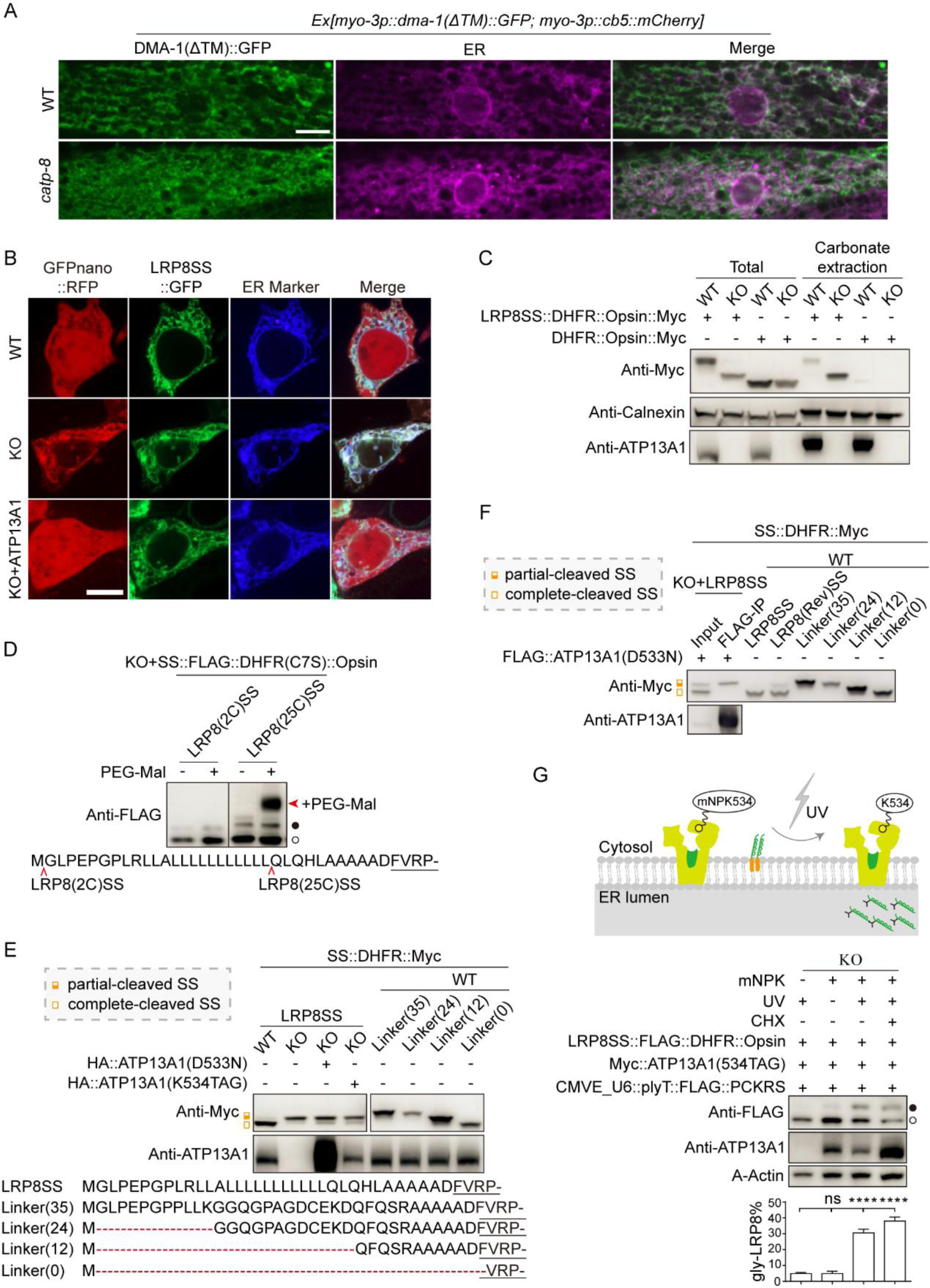
ATP13A1 facilitates translocation of cleaved P5A-dependent SS to the ER membrane. (A) Subcellular localization of DMA-1ΔTM::GFP in WT and *catp-8* worms. ER is labelled with *cb5(cytb-5.1)::mCherry*. Scale bar, 5 μm. (B) Subcellular localization of GFPnano::RFP co-expressed with LRP8SS::sfGFP in HEK293FT cells of WT, *ATP13A1* KO, or KO rescued with ATP13A1. ER is labeled with TagBFP::SEC61β. Scale bar, 5 μm. (C) Carbonate extraction of LRP8SS::DHFR::Opsin::Myc or DHFR::Opsin::Myc in WT or *ATP13A1* KO cells. Calnexin is a type I integral ER membrane protein and serves as a control. (D) PEG-maleimide modification of LRP8SS. A cysteine was inserted into LRP8SS at either the N-terminus as LRP8(2C)SS or the C-terminus as LRP8(25C)SS to investigate the integrity of LRP8SS. The cysteine in DHFR was mutated to serine (C7S) to reduce background noise. (E) LRP8SS is partially cleaved in *ATP13A1* KO cells, and cannot be rescued by ATP13A1(D533N) or ATP13A1(K534TAG). LRP8SS::DHFR::Myc and Linker::DHFR::Myc variants were transfected into WT or *ATP13A1* KO cells and blotted. Opsin tag was removed to compare lengths resulted by the differently processed SSs. LRP8SS in WT cells was translocated to the ER and thus completely cleaved. In contrast, LRP8SS in KO cells failed to translocate and was partially cleaved. (F) ATP13A1(D533N) co-immunoprecipitated with partial-cleaved LRP8SS. (G) Restoration of ATP13A1 activity by UV illumination stimulates ER translocation of mislocalized LRP8SS. A photocaged amino acid mNPK, which was genetically incorporated at position 534 by an amber suppressor tRNA/Lysyl-tRNA synthetase (PCKRS) pair to cage ATP13A1 in an inactive state. The protein synthesis inhibitor CHX was added 2hrs before restoring ATP13A1 activity by turning mNPK to lysine upon UV irradiation. Black and clear circles indicate the N-glycosylated and unglycosylated proteins, respectively. Quantification of glycosylated proteins are from 3 biological replicates and are presented as means ± SEMs; ns, not significant (*p* > 0.05); \*\*\*\**p* < 0.0001 (Student’s *t*-test). See also Figure S3.

SPP has been demonstrated to cleave mislocalized TA proteins on the ER for ERAD (McKenna et al., 2022). To assess whether LRP8SS in the incorrect topology remains intact or is cleaved in *ATP13A1* KO cells, we introduced a cysteine residue, modifiable by PEG-maleimide (Chitwood and Hegde, 2020), at either position 2 (referred to as LRP8(2C)SS) or position 25 (referred to as LRP8(25C)SS) (Figure S3A). Interestingly, the molecular weight of LRP8(25C)SS, but not LRP8(2C)SS, increased after treatment with PEG-maleimide (Figure 3D), indicating the absence of the N-terminus of LRP8SS. By comparing the length of the remaining C-terminal LRP8SS (referred to as CLRP8SS) with linkers of varying lengths (0, 12, 24, or 35 amino acids), we estimated that the length of CLRP8SS was between 12-24 amino acids (Figure 3E), further confirming that LRP8SS is partially cleaved. Despite SPP or Ste24 being quality control enzymes responsible for removing mistargeted or misfolded proteins from the ER (Ast et al., 2016; McKenna et al., 2022; Zanotti et al., 2022), the partial cleavage of LRP8SS was not inhibited by the SPP inhibitor (Z-LL)_2_- ketone or the Ste24 inhibitor lopinavir (Figure S3B). Therefore, further characterization is needed to understand the surveillance mechanism by which misoriented P5A-dependent SSs in the ER are recognized and processed into short SSs.

The ATP13A1(D533N) variant specifically co-immunoprecipitated with CLRP8SS, excluding the translocated passenger protein, as illustrated in Figures 3F and S3C. Subsequently, we explored whether CLRP8SS represents the product processed by protein quality control due to its incorrect topogenesis in *ATP13A1* KO cells or an intermediate product during P5A-mediated translocation. To address this, we introduced a light-responsive non-canonical amino acid (ncAA), methyl-o- nitropiperonyllysine (mNPK), at position K534, adjacent to the catalytic residue D533, to restrain ATP13A1 in an inactive state (Kneuttinger et al., 2019). The inactivity was confirmed in Figures 3G and S3D. Upon UV irradiation to remove the caging group from mNPK, thereby restoring ATP13A1 activity, the stuck LRP8SS translocated to the ER, as depicted in Figure 3G. Additional confirmation of the translocated LRP8SS being preexisting, rather than newly synthesized, was achieved by adding cycloheximide (CHX) 2 hours before UV illumination to inhibit protein synthesis. Despite this inhibition, reactivating photo-caged ATP13A1 increased the translocated portion of ER-stuck LRP8SS, as indicated in Figure 3G. This data strongly supports the notion that replenishing ATP13A1 can dislocate misoriented LRP8SS from the ER, thereby facilitating its translocation.

### ATP13A1 Engages GET3 for the ER translocation of P5A-dependent SS

Investigation into the fate of CLRP8SS subsequent to dislocation by ATP13A1 revealed that the ER translocation of CLRP8SSs with different estimated lengths (CLRP8SS-1, CLRP8SS-2, and CLRP8SS-3) no longer required ATP13A1, as depicted in Figure 4A. This suggests the involvement of other factors in the ER translocation of CLRP8SS after dislocation by ATP13A1. Comparative analysis of the proteins interacting with LRP8SS in WT cells, PAT-3SS in WT, or LRP8SS in *ATP13A1* KO cells through mass spectrometry revealed specific enrichment of GET3 by LRP8SS in WT cells (Figure S4A). The GET complex, consisting of the targeting factor GET3 and its ER receptor GET1/2 (WRB/CAML), is a well-established insertase for TA-proteins inserting into the ER membrane (Guna et al., 2018; Mariappan et al., 2011; Stefanovic and Hegde, 2007). Using the ATPase-deficient mutant GET3(D74N), which constitutively binds to TA proteins, and the GET3(I193D) mutant, defective in binding to TA-proteins(Mateja et al., 2009), we validated the interaction between GET3 and CLRP8SS. The direct interaction was disrupted by GET3(I193D) but not GET3(D74N), as shown in Figure S4D. Furthermore, incorporation of pAzF into the substrate-binding hydrophobic groove of GET3, revealed by its crystal structure (Mateja et al., 2015), demonstrated that LRP8SS contacted with the residues M146 and A149 in the hydrophobic groove of GET3 (Figures S4E and S4F).

**Figure 4.**
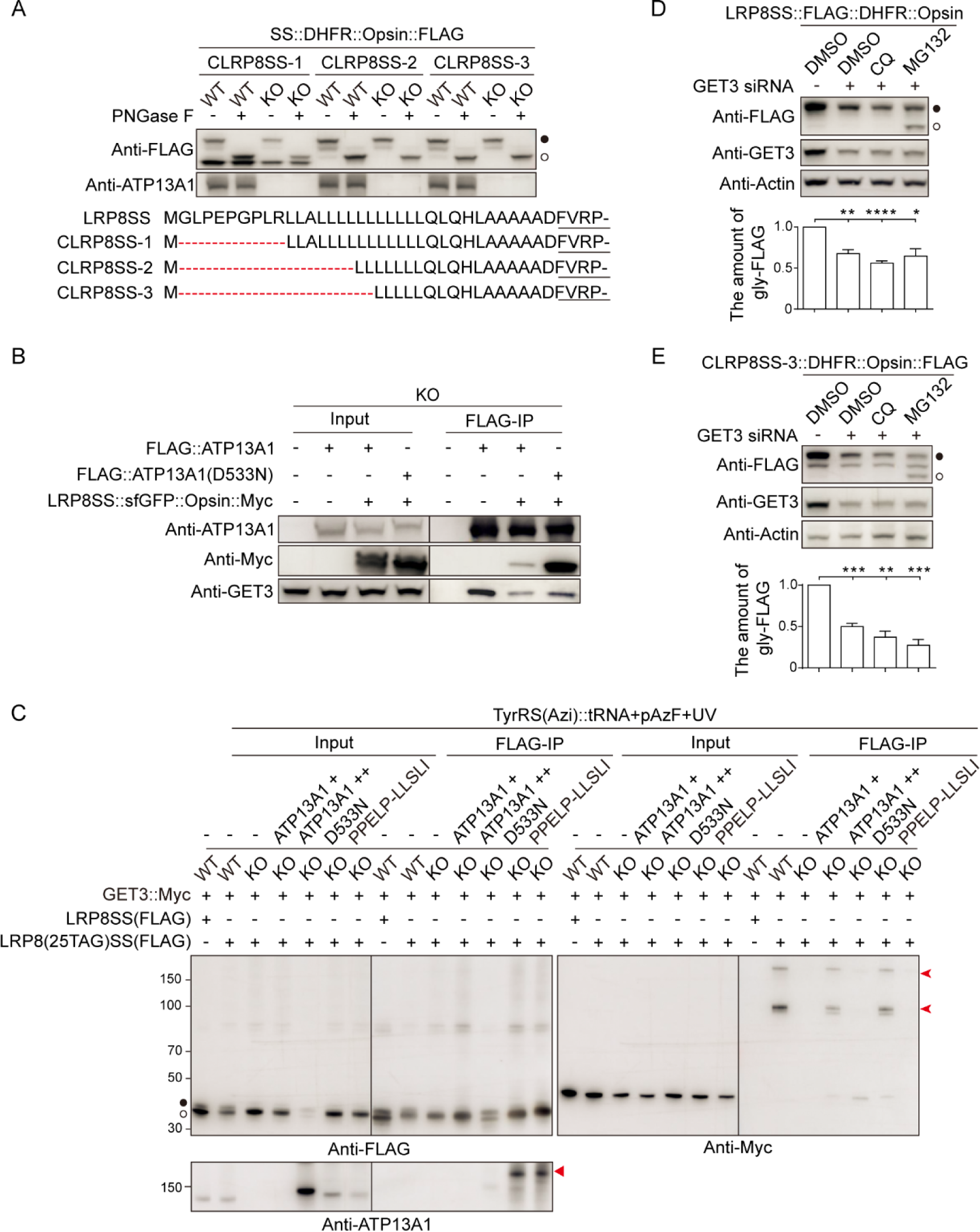
ATP13A1 engages GET3 for ER translocation of P5A-dependent SSs. (A) ATP13A1 is dispensable for the ER translocation of CLRP8SS variants. LRP8SS, CLRP8SS-1, CLRP8SS-2, and CLRP8SS-3 fused with DHFR::Opsin::FLAG was transfected into WT and *ATP13A1* KO cells, and then lysed and treated with or without PNGase F. (B) ATP13A1 or ATP13A1(D533N) co-immunoprecipitated with GET3 and LRP8SS. (C) PPELP motif of ATP13A1 is essential for the interaction between GET3 and LRP8SS. LRP8(25TAG)SS::sfGFP::Opsin::FLAG or LRP8SS::sfGFP::Opsin::FLAG was co-transfected with ATP13A1 variants in WT or *ATP13A1* KO cells. + or ++ indicates low or high expression of the ATP13A1 variants by the TC573 or pEF6a vector, respectively. The red arrow represents the band of GET3 crosslinked with LRP8SS. (D-E) GET3 is required for the ER translocation of LRP8SS and CLRP8SS. Control or GET3 siRNA was co-transfected with LRP8SS::FLAG::DHFR::Opsin (D) and CLRP8SS-3::DHFR::Opsin::FLAG (E) into WT cells treated with DMSO, chloroqine, or MG132. Black and clear circles indicate the N-glycosylated and unglycosylated proteins, respectively. See also Figures S4 and S5.

LRP8SS binding to GET3 was significantly disrupted in *ATP13A1* KO cells (Figure 4C). Furthermore, an interaction between ATP13A1 and GET3 was observed (Figure 4B). Thus, we hypothesized that ATP13A1 facilitates the displacement of CLRP8SS from the ER to GET3. Cryo-EM structures revealed that the transport domain of Spf1 is constituted by TM1-TM6, with TM4 being notable forbreaking into TM4a and TM4b at the conserved PPELP motif. During the E1P-to-E2P transition of ATP13A1, TM4a and TM4b undergo a substantial conformational change, leading the substrate-binding pocket to shift from the inward-open (apo) to the outward-open states (McKenna et al., 2020).

To assess the significance of the PPELP motif in ATP13A1 for delivering LRP8SS to GET3, we introduced a mutation (PPELP-LLSLI). Similar to the hydrolytic mutant D533N, LRP8SS translocation was abolished in the PPELP-LLSLI mutant (Figure S4C). Despite ATP13A1 variants PPELP-LLSLI and D533N retaining the ability to bind LRP8SS (Figures S5B and S5C) and GET3 (Figure S5A), PPELP- LLSLI disrupted the interaction between GET3 and LRP8SS (Figure 4C). Thus, our results indicate that ATP13A1 plays a crucial role in displacing LRP8SS to GET3. We subsequently investigated the requirement of GET3 for the ER translocation of P5A- dependent SSs. Indeed, siRNA-mediated depletion of GET3 significantly diminished the ER translocation of both LRP8SS and CLRP8SS (Figures 4D and 4E). Moreover, the untranslocated LRP8SS and CLRP8SS, generated by GET3 knockdown, were stabilized by the proteasome inhibitor MG132 (Figures 4D and 4E), suggesting their instability in the cytosol and subsequent ERAD. Our findings collectively support a model wherein ATP13A1 collaborates with GET3 for the ER translocation of CLRP8SS.

In addition to their role as ER targeting factors, GET3 functions as chaperones to sequester TA protein aggregation (Powis et al., 2013). Given GET3’s downstream position relative to ATP13A1, we explored whether GET3 acts as a target factor or molecular chaperone in facilitating the ER translocation of P5A-dependent SSs. We examined the interaction between LRP8 truncations of varying lengths and GET3. While the length of passenger proteins was critical for LRP8 translocation, it did not affect the interaction between LRP8 and GET3 (Figures S5D and S5E). This observation suggests that GET3 more likely functions as a target factor in LRP8 translocation.

### P5A-dependent SS Directed to the ER through Interaction with the SRP Complex

Following the dislocation of LRP8SS from the ER by ATP13A1, we sought to understand the immediate ER targeting mechanism for LRP8SS post-translation. The known pathways for SS-mediated ER targeting involve SRP-dependent co- translational or the Sec62-Sec63 dependent post-translational translocation pathway (Figure 5A). Surprisingly, despite P5A-dependent SSs being post-translationally translocated (Figure 3G), a *sec62* mutation showed no impact on DMA-1SS secretion from the intestine to the pseudocoelom in worms (Figure 5B).

**Figure 5.**
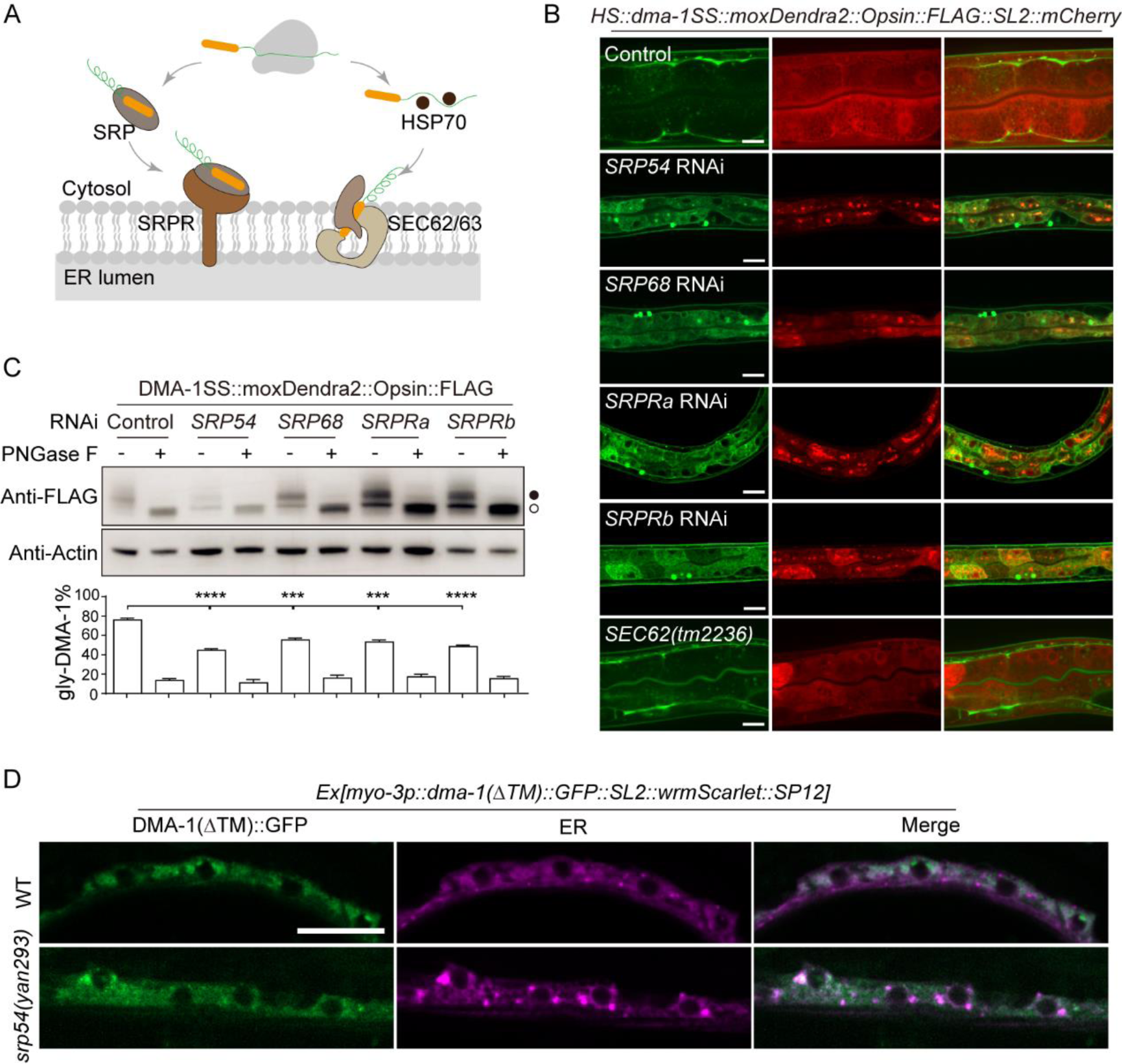
P5A-dependent SS targets ER via the SRP complex. (A) Diagram illustrating SS-mediated ER targeting via signal recognition particle (SRP)-dependent co-translational or Sec62-Sec63-dependent post-translational translocation pathway. (B) Secretion of DMA-1SS::moxDendra2::Opsin::FLAG reporter in worms treated with RNAi in control (L4440), *SRP54*, *SRP68*, *SRPR*α, *SRPR*β, or *sec62 mutants*. Scale bar, 10 μm. (C) Western blot of DMA-1SS::moxDendra2::Opsin::FLAG reporters in (B). Black and clear circles indicate the N-glycosylated and the unglycosylated proteins, respectively. Quantifications are from 3 biological replicates and presented as means ± SEMs; \*\*\**p* < 0.001; \*\*\*\**p* < 0.0001 (Student’s *t*-test). (D) Subcellular localization of DMA-1ΔTM::GFP in WT and *srp54* mutant worms. ER is labelled by SP12::mCherry. Scale bar, 5 μm. See also Figure S6.

To identify factors crucial for ER targeting, a forward genetic screen utilizing DMA-1SS::moxDendra2::Opsin::FLAG reporter revealed a dominant-negative mutation, G257R, in SRP54 severely disrupting DMA-1SS secretion in worms (Figure S6A). Depletion of SRP complex components (SRP54, SRP68, and SRPR) through RNA interference resulted in the retention and unglycosylation of DMA-1SS in the intestine (Figures 5B and 5C). The G257R mutation in *srp54(yan293)* impeded the ER targeting of DMA-1ΔTM::GFP, causing it to disperse in the cytosol (Figure 5D). In line with the worm observations, overexpression of G257R, as well as two disease-causing mutations in SRP54, T115A and T117del (Juaire et al., 2021), reduced LRP8SS levels in HEK293FT cells (Figures S6B). This reduction is attributed to the proteasome degradation of LRP8SS (Figure S6C), as SRP mutants failed to deliver LRP8SS to the SRP receptor, SRPR. Moreover, SRPR co- immunoprecipitated with ATP13A1 (Figures S6D-S6F). In conclusion, these findings indicate that P5A-dependent SS is initially targeted to the ER via the SRP complex, and ATP13A1 plays a central role in the translocation by bridging the SRP-mediated initial targeting and the GET3-mediated second targeting.

### Translocation of P5A-dependent SS to the ER Requires the SEC61 Translocon

We investigated the involvement of SEC61 translocon or EMC insertase in the ER translocation of P5A-dependent SSs. Knocking down the α or γ subunit of SEC61 disrupted the glycosylation and secretion of DMA-1SS (Figures 6A-6C, S7A-S7B), while *emc-6* deficiency did not affect DMA-SS secretion (Figure S7C). These results suggest that SEC61 translocon, but not the EMC complex, is engaged in the ER translocation of P5A-dependent SSs. Further investigation revealed an interaction between A TP13A1 and endogenous SEC61 (Figures 6D), and the catalytic mutation D533N disrupted this interaction (Figures 6D and 6E). Notably, ATP13A1(D533N) not only co-immunoprecipitated with more LRP8SS (Figure 6F) but also enhanced the interaction between LRP8SS and SEC61 (Figures 6D). This suggests that the ATP13A1 mutation leads to more SSs trapped in both ATP13A1 and SEC61. Conversely, overexpressing ATP13A1 enhanced the translocation efficiency of LRP8SS, consequently reducing the interaction between LRP8SS and SEC61 (Figure 6F). In summary, our results support the idea that SEC61 cooperates with ATP13A1 for LRP8SS translocation.

**Figure 6.**
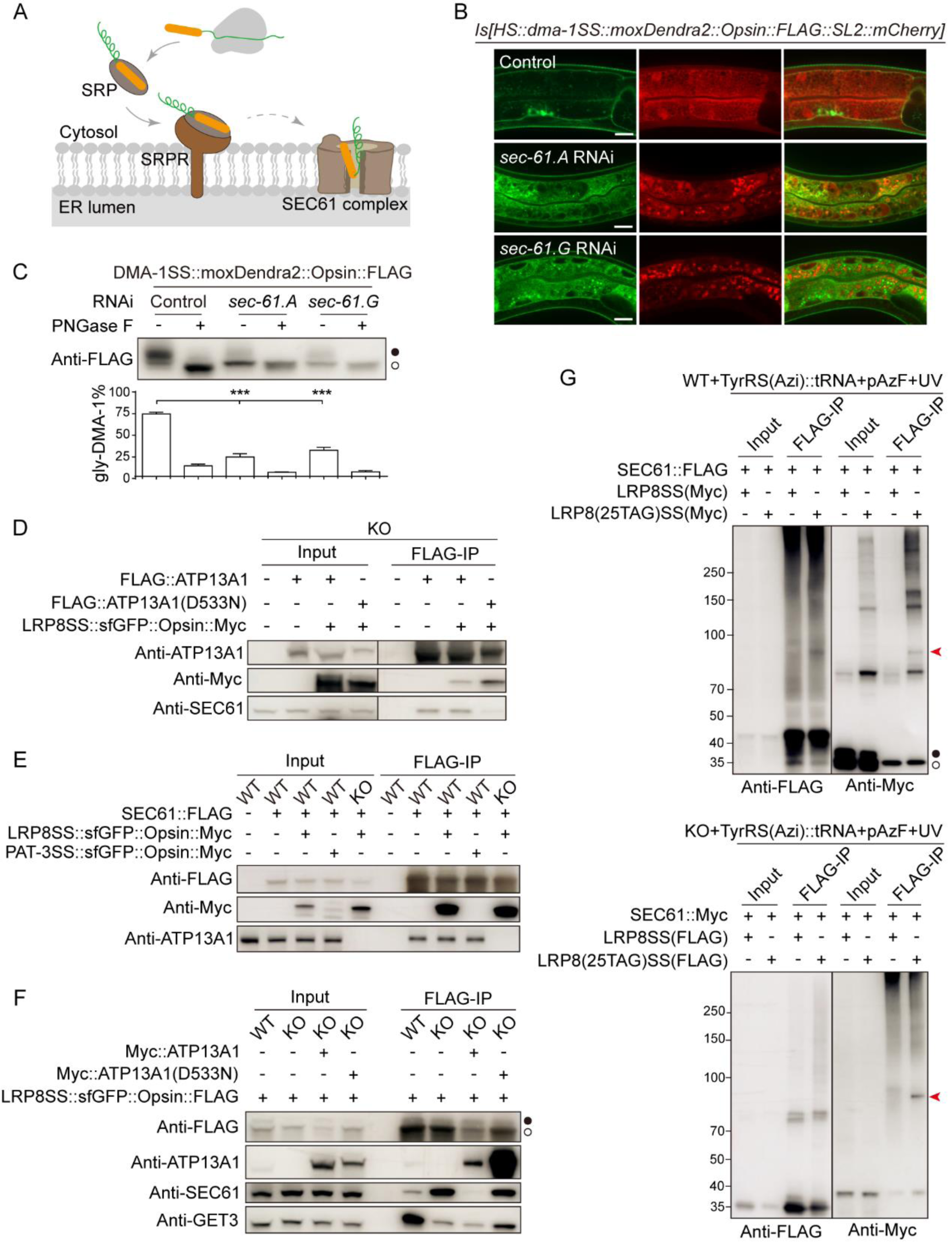
ER translocation of P5A-dependent SS requires SEC61 translocon. (A) Diagram illustrating SS-mediated ER translocation via SEC61. (B) Secretion of DMA-1SS::moxDendra2::Opsin::FLAG reporter in worms treated with RNAi in control (L4440), *sec-61.A*/*SEC61*α, and *sec-61.G/SEC61*γ. Scale bar, 10 μm. (C) Western blot of DMA-1SS::moxDendra2::Opsin::FLAG reporters in (B). Quantifications are from 3 biological replicates and presented as means ± SEMs; \*\*\**p* < 0.001 (Student’s *t*-test). (D) Co-immunoprecipitation of ATP13A1 variants, LRP8SS, and endogenous SEC61. (E) Co-immunoprecipitation of SEC61, LRP8SS, and endogenous ATP13A1. (F) Co-immunoprecipitation of ATP13A1 variants, LRP8SS, and endogenous SEC61 or GET3. (G) LRP8SS is crosslinked with SEC61 in WT or *ATP13A1* KO cell. The red arrow represents the SEC61 band crosslinked with LRP8SS. Black and clear circles indicate the N-glycosylated and the unglycosylated proteins, respectively. See also Figure S7.

## Discussion

In this study, we demonstrate that P5A-dependent SSs exhibit an increased error- prone nature during topogenesis when directed to the ER through the SRP complex. The misoriented SSs undergo processing and dislocation from the ER membrane by ATP13A1, subsequently engaging the GET3 complex for a second chance of ER targeting, followed by translocation through the SEC61 translocon (Figure 7).

**Figure 7.**
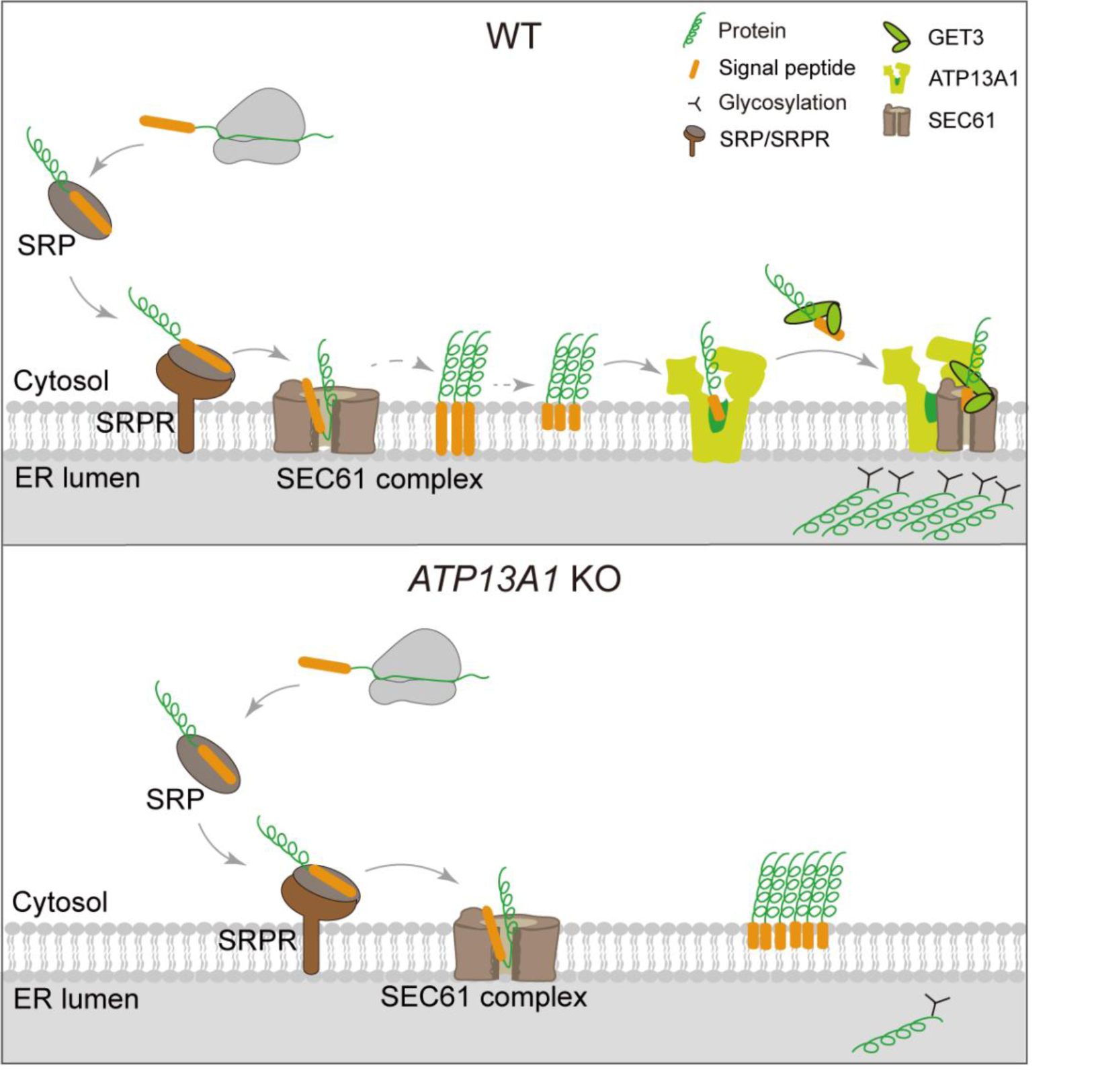
ATP13A1 dislocates cleaved SSs to GET3 and engages SEC61 for ER translocation. Atypical SSs with high hydrophobicity and lack of characteristic charges are incorrectly targeted into the ER through the SRP complex and are subjected to partial cleavage. ATP13A1 dislocates cleaved SSs to the targeting factor GET3 and engages SEC61 for ER translocation in WT cells. However, in *ATP13A1* KO cells, the majority of atypical SSs are targeted to the ER in an incorrect topology, while only a small portion of atypical SSs are correctly targeted and translocated through SEC61.

SSs play crucial roles in protein biogenesis and trafficking by interacting with the engaging SRP complex for ER targeting and ensuring accurate topogenesis during protein translocation. A typical SS comprises three components: an N-terminal region containing positively charged residues facilitating binding to the SRP, adopting the N^cyto^ topology by interacting with lipids in the ER membrane at the cytosolic side; a hydrophobic core in the central region for insertion into the ER membrane; and a C terminus housing the SPP cleavage site, removed post-import. Despite their importance, SS sequences exhibit high diversity and degeneracy, posing a challenge in deciphering the molecular codes governing their interpretation. The translocon- associated protein (TRAP) complex and the translocating chain-associated membrane protein (TRAM) are known to initiate co-translational translocation of weakly hydrophobic SSs through Sec61 (Fons et al., 2003; Voigt et al., 1996). Similarly, the Sec62-Sec63 complex facilitates the post-translational translocation of weakly hydrophobic SSs (Schorr et al., 2020). Our findings reveal that P5A-dependent SSs deviate from the typical pattern, displaying unusually high hydrophobicity in the central core and a lack of positive charges at the N-terminus (Figures S1A and S1B), resulting in misoriented insertion into the ER membrane (Figures 2A and 3). These misoriented P5A-dependent SSs undergo partial cleavage, recognized and dislocated by ATP13A1 from the ER (Figure 3). The cryo-EM structure indicates that the potential substrate-binding pocket of SPF1, the yeast ortholog of P5A ATPase, is large and shallow, alternating its orientation towards the ER lumen or cytosol while maintaining lateral openings accessibility (McKenna et al., 2020). Consequently, the partial cleavage of SSs in misoriented proteins may contribute to substrate selection and render them biophysically more accessible for dislocation from the ER by ATP13A1.

While GET3 is widely recognized as a targeting factor for the insertion of TA proteins to the ER, our findings demonstrate that ATP13A1 collaborates with GET3 to direct dislocated SSs to the ER. This offers a secondary opportunity for ER translocation by the SEC61 complex, thereby preventing their degradation by the proteasome (Figure 4). Notably, both the PPELP-LLSLI and D533N mutations of ATP13A1 impede SS translocation, yet they retain the ability to interact with GET3 (Figure 4). However, only the PPELP-LLSLI mutation disrupts the interaction between GET3 and SS (Figure 4). The most plausible model to elucidate these results suggests that conformational changes and ATP hydrolysis of ATP13A1 are necessary for dislocating SSs to GET3 and subsequently releasing GET3-SSs, respectively. Without conformational change, ATP13A1 cannot pass SSs to GET3. Without ATP hydrolysis, the ATP13A1-GET3-SS complex is likely arrested, impeding further translocation. This is supported by the observation that the D533N mutation abolishes the ATP13A1-SEC61 interaction (Figure 6D) but enhances interactions between SSs and ATP13A1 (Figure 6F). The interaction and collaboration between SEC61 and ATP13A1 for translocating partially cleaved SSs require further investigation. Furthermore, our study aligns with McKenna et al.’s findings that ATP13A1 is crucial for dislocating mislocalized mitochondrial TA proteins from the ER (McKenna et al., 2022; McKenna et al., 2020). It remains to be established whether GET3 or distinct factors cooperate with ATP13A1 in the mitochondrial insertion of dislocated TA proteins.

In summary, our investigation delineates a comprehensive translocation pathway crucial for error-prone SSs, with ATP13A1 playing a pivotal role in dislocating misoriented SSs and facilitating subsequent translocation by engaging the targeting factor GET3 and the Sec61 translocon. Evolutionarily, the ER-targeting SSs in eukaryotes possess less positively charged N-termini compared to their plasma- membrane targeting counterparts in prokaryotes, a feature evolved to avoid cross-talk with targeting sequences destined for bacterial-derived mitochondria (Garg and Gould, 2016). The emergence of P5A-ATPase in eukaryotes likely addresses this need by coordinating the protein quality control system and the translocation machinery. Consequently, our study provides valuable insights into how the components of a secretory apparatus act in a substrate-dependent manner to maintain high efficiency and fidelity.

## Methods

### RESOURCE AVAILABILITY

#### Lead Contact

Further information and requests for resources and reagents should be directed to and will be fulfilled by the Lead Contact, Yan Zou.

#### Materials Availability

All *C. elegans* strains and plasimds generated in this study are available on request from the Lead Contact.

#### Data and Code Availability

Proteomics data have been deposited at MassIVE and are publicly available as of the date of publication. Accession numbers are listed in the key resources table.

This paper does not report original code.

Any additional information required to reanalyze the data reported in this paper is available from the lead contact upon request.

### EXPERIMENTAL MODEL AND SUBJECT DETAILS

#### Animals

N2 Bristol was used as the wild-type strain. *C. elegans* strains were grown on nematode growth medium (NGM) plates seeded with OP50 *E. coli*, and maintained at 20°C. Details and a complete list of strains in this study are shown in Table S1.

#### Cell lines

All cell lines were maintained in DMEM with 4.5 g/L D-Glucose, L-Glutamin supplemented with 10% fetal bovine serum (NSERA, S711-001S) at 37 °C and 5% CO_2_. HEK293FT *ATP13A1* KO cell was obtained by knocking out *ATP13A1* in wild-type HEK293FT using Crispr-Cas9 technology, which was used in our previous article (Feng et al., 2020; Li et al., 2021).

### METHOD DETAILS

#### Molecular Biology and Transgenesis

Reagents used and generated in this study, see Key Resources Table. Most of the plasmid constructs were generated in pSM/pEGFP.C1/pEF6a/pRK5 vector backbone. Germline transformation of *C. elegans* was performed using standard techniques (Mello and Fire, 1995). *pCFJ104, pCFJ90, odr- 1p::gfp, or odr-1p::rfp* plasmid was injected at 2.5-60 ng/µL as co-injection marker.

#### PNGaseF Treatment in Mammalian Cells

PNGaseF treatment was performed according to a previously reported protocol with modification (Feng et al., 2020). Cells were lysed on ice for 30 min with 150 μL of NP40 lysis buffer (50 mM Tris-HCl, pH 7.4, 150 mM NaCl, 1 mM EDTA, 0.5% NP40) containing 1X Protease Inhibitor Cocktail and centrifugated at 15,000 rpm for 10 min at 4 °C. The supernatant was divided into two groups and treated with PNGase F (a gift from Dr. Yu Cao) and ddH_2_O for 1 hr at room temperature, respectively. After the treatment with PNGase F, the samples were added with 5X Sample buffer (250 mM Tris-HCl pH 6.8, 5% SDS, 50% glycerol, 500 mM DTT) to a final concentration at 1X, incubated at 65 °C for 10 min, and analyzed by immunoblotting with indicated antibodies in Figures 1F and 4A. The PNGase F treatment in *C. elegans* can be seen as “Knock down of SEC61 complex or SRP/SRPR complex” (for images showed in Figures 1C, 5C and 6C).

#### (Z-LL)2-ketone and Lopinavir Treatment of HEK293 *ATP13A1* KO Cells

(Z-LL)_2_-ketone and lopinavir treatment was performed according to a previously reported protocol with modification (Ast et al., 2016; Weihofen et al., 2000). Briefly, 3.2 x 10^5^ HEK293FT *ATP13A1* KO cells from six 3.5 cm plates were transfected with LRP8::DHFR::Myc and then aliquoted for six treatments (#1-#6). Six hours after transfection, ddH2O, 10 μM MG132, 50 μM (Z-LL)2-ketone, 50 μM (Z- LL)_2_-ketone with 10 μM MG132, 20 μM lopinavir, and 20 μM lopinavir with 10 μM MG132 (final concentrations) were added to #1-#6 for an additional 15 hrs, respectively. Then the cells were pelleted and extracted by NP40 lysis buffer containing 1X Protease Inhibitor Cocktail on ice for 30 min. The soluble fraction of the cell lysates was isolated via centrifugation at 12,000 rpm for 10 min at 4 °C. The supernatants were analyzed by immunoblotting with indicated antibodies in Figure S3B.

#### siRNA-Mediated Knock Downs, CQ and MG132 Treatments in Cells

1.2 x 10^5^ HEK293FT cells from four 3.5 cm plates were cotransfected using Lipofectamine 2000 (Invitrogen) according to Figures 4D and 4E and then aliquoted for four treatments (#1-#4). #1 is the control group, and #2-#4 are the knockdown groups. The cells were firstly co-transfected with siRNAs and the reporter, and were transfected again with the siRNAs 24 hrs later to enhance the knockdown efficiency (Figure 4D and 4E). 12 hours after the second transfection, MG132 (10 μM final concentration) and chloroquine (CQ, 50 μM final concentration) were added to the cell culture medium for an additional 15 hrs and 12 hrs, respectively. After the treatment with MG132 and CQ, the cell lysates were extracted by NP40 lysis buffer containing 1X Protease Inhibitor Cocktail for 30 min on ice. The soluble fraction of the cell lysates was isolated via centrifugation at 15,000 rpm in a microcentrifuge for 10 min at 4 °C. The supernatants were analyzed by immunoblotting with indicated antibodies in Figures 4D and 4E. The treatment of CQ and MG132 in Figures S6C followed similar protocols.

#### Knock Down of SEC61 Complex or SRP/SRPR Complex

RNAi feeding experiments were conducted following the protocol (Kamath et al., 2001). An appropriate amount of *C. elegans* were cultured to day 2 adult, and synchronized to obtain eggs. The eggs were added onto NGM agar plates with control, *sec-61.A* or *sec-61.G* RNAi bacteria. After incubated at 20 °C for 2 days, the grown up worms were treated at 33 °C for 60 min to induce the expression of the DMA- 1SS or PAT3-SS reporter, and then collected after incubation at room temperature for 6 hrs.

Synchronized L4 stage nematodes were washed with 0.01% Triton/M9 solution for 3-5 times, and then added onto the NGM plates with control (about 8 nematodes per dish) or SRP/SRPR RNAi bacteria (about 50 nematodes per dish). After incubated at 20 °C for 2.5 days, the worms were treated at 33 °C for 60 min, and then collected after incubation at room temperature for 6 hrs.

The harvested worms were resuspended with Sample buffer and lysed at 95 °C for 3 min. After centrifuged at room temperature at 15,000 rpm for 5 min, the supernatant was transferred to a new EP tube, and added with 10% NP40 (Solarbio) to a final concentration of 1%. Then the supernatant was divided into two groups and treated with PNGase F and ddH_2_O for 3 hours at 37 °C, respectively. After the treatment, the samples were added with 0.1% bromophenol blue and analyzed by immunoblotting with indicated antibodies in Figures 5C, 6C, S7A and S7B.

#### Carbonate Extraction

The separation of membrane integral proteins by carbonate extraction was performed according to a previously reported protocol with modification (Chitwood, 2020; Fujiki et al., 1982). Briefly, 2.4 x 10^6^ WT or 3.6 x 10^6^ *ATP13A1* KO cells from two 6 cm plates were cotransfected according to Figures 3C. The next day, cells were harvested, washed twice with PBS, resuspended with 200 μL RM buffer (10 mM HEPES, pH 7.4, 250 mM Sucrose, 2 mM MgCl2, 1X Protease Inhibitor Cocktail), lysed with Bioruptor®Plus high-power mode for 5 cycles (sonication cycle: 30 s “ON”, 30 s “OFF”), and then centrifuged at 600 x g for 3 min to remove nuclei. The supernatant was transferred to a new EP tube and centrifuged at 15,000 x g for 5 min. Then 150 μL of supernatant was transferred to an ultra- centrifuge tube, mixed well with 750 μL of 250 mM Sucrose and 100 μL of 1 M Na_2_CO_3_, and incubated on ice for 30 min. The mixture was centrifuged at 55,000 rpm for 1 hr in Optima™ MAX-TL Ultracentrifuge (BECKMAN COULTER). Then the pellet was dissolved with 200 μL 1X Sample buffer, and incubated at 95 °C for 10 min before immunoblotting analysis (for images showed in Figures 3C).

#### Detecting the Integrity of Signal Sequence by PEG-Mal

The PEG-Mal treatment was performed according to a previously reported protocol with modifications. 8 x 10^5^ HEK293FT *ATP13A1* KO cells on 3.5 cm plates were transfected with LRP8(2C)SS::FLAG::DHFR(C7S)::Opsin and LRP8(25C)SS::FLAG::DHFR(C7S)::Opsin, respectively.

24 hours after transfection, the cells were collected and extracted by lysis buffer (50 mM Tris-HCl, pH 7.4, 150 mM NaCl, 1mM EDTA, 1% Triton X-100, 1X Protease Inhibitor Cocktail) for 10 min on ice. The soluble fraction of the cell lysates was isolated via centrifugation at 15,000 rpm for 10 min at 4 °C. The supernatant was immunoprecipitated with Anti-FLAG M2 affinity Gel (Sigma). Samples were eluted off from the beads via 50 µL eluent (50 mM HEPES pH 7.4, 5% SDS). 19 µL of elute was mixed with 1 µL of 20 mM PEG-Mal (Sigma-Aldrich) or 1 µL ddH_2_O, and incubated at room temperature for 1.5 hrs in dark room. Immunoblotting was performed after the samples added with 100 µL 1X Sample buffer and treated at 65 °C for 10 minutes (for images showed in Figures 3D).

#### Photo-activation of ATP13A1

Photo-control was performed according to a previously reported protocol with modifications (Gautier et al., 2010; Kneuttinger et al., 2019). Briefly, four groups of 8 x 10^5^ HEK293FT *ATP13A1* KO cells on 3.5 cm plates were cotransfected with 0.5 µg pEGFP.C1_LRP8::FLAG::DHFR::Opsin, 1 µg pRK5_Myc::ATP13A1(K534TAG), and 1 µg pEGFP.C1_4x (CMVE_U6::plyTtRNA)_EF1a::FLAG::PCKRS and then aliquoted for four treatments (#1-#4). Six hours after transfection, the medium was changed to DMEM containing 10% FBS and 1 mM Methyl-o- nitropiperonyllysine (mNPK, Sigma-Aldrich) for #2-#4, and changed to DMEM containing 10% FBS and NaOH (a control solution for mNPK) for #1. The next day, the protein synthesis inhibitor cycloheximide (CHX, 50 µg/mL) and DMSO was added to the #4 and #1-#3, respectively. 2 hours after treatments, the medium was changed to 1 mL PBS, after wrapping #2 in aluminum foil but removing the lid of the #1, #3-#4, photo-control by exposure to 365 nm UV light with a UV lamp (SCIENTZ 03-II) in a dark room for 1 min. PBS was exchanged for 2 mL of DMEM containing 10% FBS for #1-#3, and exchanged for 2 mL of DMEM containing 10% FBS and 50 µg/mL CHX for #4. Cells were collected after 6 hours of re-culture at 37°C, 5% CO_2_. The cell lysates were extracted by NP40 buffer containing 1X Protease Inhibitor Cocktail for 30 min on ice. The soluble fraction of the cell lysates was isolated via centrifugation at 15,000 rpm in a centrifuge for 10 min at 4 °C. The supernatants were analyzed by immunoblotting with indicated antibodies in Figures 3G.

#### Site-Specific Photo-Crosslinking

Photo-crosslinking was performed according to previously reported protocols with modifications (Naganathan et al., 2013; Zhang et al., 2021). Note that the whole experiment needs to be protected from light. Briefly, 3.5 x 10^6^ HEK293FT cells or 4 x 10^6^ HEK293FT *ATP13A1* KO cells on 10 cm plates were cotransfected with 4 µg of plasmids containing the amber stop codon at the desired position, 4 µg TyrRS(Azi)::4x(U6-tRNA), and 4 µg related other plasmids as indicated in Figures 2D and 2F, 4C, 6G, S4D and S4F, and S5C and S5D. Six hours after transfection, the medium was changed to DMEM containing 10% FBS and 1 mM 4-Azido-L-phenylalanine (pAzF, MedChemExpress). The next day, the cells were collected, washed 3 times with PBS, and then re-suspended with 1 mL of PBS, and photocrosslinked by exposure to 365 nm UV light with a UV lamp (SCIENTZ 03-II) in a dark room on ice for 30 min. Then the cells were collected by centrifugation at 1,000 x g for 3 min, denatured and co-immunoprecipitated, and analyzed by western blotting (for images showed in Figures 2D and 2F, 4C, 6G, S4D and S4F, and S5C and S5D).

#### Native Immunoprecipitation

Heat at 95 °C for 5 min to dissolve digitonin for later use. The cells pellet (approximately 100 μL/sample) were resuspended with 300 μL of lysis buffer (50 mM Tris-HCl, pH 7.4, 150 mM NaCl) containing 1X Protease Inhibitor Cocktail and 1% digitonin and incubate on ice for 2 hrs. The samples were added with 1 mL lysis buffer and centrifuged at 15,000 rpm at 4 °C for 10 min, and then transfer the supernatant to a new EP tube and mix well. Mix 40 μL of supernatant, 40 μL ddH_2_O, and 20 μL of 5X Sample buffer and incubate at 65°C for 10 min as input. For immunoprecipitations under native conditions, the remaining supernatant were added to EP tube containing ANTI-FLAG M2 Affinity Gel and incubated in a cold room for 4-6 hrs. After incubation, the samples were washed four times (10 min each) at 4 °C with 1 mL wash buffer (50 mM Tris-HCl, pH 7.4, 500 mM NaCl, 0.03% digitonin). Then transfer beads to a new EP tube, aspirate the supernatant as much as possible, add 80 μL of 1X Sample buffer and incubate at 65 °C for 10 min. The sample were analyzed by immunoblotting with indicated antibodies in Figures 2B, 3F, 4B, 6D-F, S3C, and S5A-B and S5E. The lysate used in Figure S6D-F is NP40 buffer and the Wash buffer is 0.1% NP40 buffer (150 mM NaCl, 0.1% NP40 and 50 mM Tris-HCl, pH 7.4), other steps followed similar protocols.

#### Denatured-Reduced Immunoprecipitation

Unless otherwise indicated, all photo-crosslinked samples were referred to denatured-reduced co-immunoprecipitation modified from a previous reported protocol (Chitwood and Hegde, 2020). The cells pellet (approximately 100 μL/sample) were resuspended with 200 μL lysis buffer (50 mM Tris-HCl, pH 7.4, 150 mM NaCl) containing 1X Protease Inhibitor Cocktail and disrupted using Bioruptor®Plus high-power mode for 5 cycles (sonication cycle: 30 s “ON”, 30 s “OFF”). The samples were centrifuged at 1,000 x g at 4 °C for 5 min to remove nuclei, and then transfer the supernatant to a new EP tube and mix well. Mix 20 μL of supernatant, 60 μL ddH_2_O, and 20 μL of 5X Sample buffer and incubate at 95°C for 10 min as input. For immunoprecipitations under denaturing-reducing conditions, the remaining supernatant were first denatured-reduced in lysis buffer containing 1% SDS and 10 mM DTT for 10 min at room temperature and boiled for 10-15 min. After SDS and DTT treatment, DTT was quenched by 30 mM N-Ethylmaleimide (NEM, MedChemExpress) for 20-30 min at room temperature. Samples were dilute 10-fold with lysis buffer containing 1% TritonX-100, and then immunoprecipitated in batch with desired antibodies at 4 °C rotating end-over-end for 4-6 hrs. After incubation, the samples were washed 4 times (10 min each) at 4 °C with 5 mL of wash buffer (50 mM Tris-HCl, pH 7.4, 500 mM NaCl, 1% TritonX-100). Then transfer beads to a new 1.5 mL EP tube, aspirate the supernatant as much as possible, add 80 μL 1X Sample buffer and incubate at 95 °C for 10 min (for images showed in Figures 2D, 2F, 4C, 6G, S4D and S4F, and S5C and S5D).

#### The Pup-IT Labeling Assay

Pup-IT labeling assay were performed as described (Liu et al., 2018). Briefly, three 1.2 x 10^6^ HEK293FT cells on 6 cm plates were cotransfected according to Figures S2B. Six hours after transfection, the medium was changed to DMEM containing 10% FBS and 4 mM Biotin. The next day, the cells were collected and lysed on ice for 30 minutes with NP40 buffer containing 1X Protease Inhibitor Cocktail. After incubation, the cells were centrifuged at 15,000 x g for 10 min and the supernatant was transferred to 1.5 mL EP tube containing Anti-FLAG M2 affinity Gel and then incubate in a cold room for 4 hrs. The samples were washed four times (10 min each) at 4 °C with 1 mL of Wash buffer. Then transfer beads to a new EP tube, aspirate the supernatant as much as possible, add 80 μL of 1X Sample buffer and incubate at 65 °C for 10 min. The sample were analyzed by immunoblotting with indicated antibodies in Figures S2B.

#### Signal Sequence Prediction and Hydrophobicity Calculation

Signal sequences were predicted by SignalP 5.0 (https://services.healthtech.dtu.dk/services/SignalP-5.0/) (Armenteros et al., 2019)(for images showed in Figure S1A and S2B). The hydrophobic core regions in signal sequences were predicted by Phobius (https://phobius.sbc.su.se/) (Kall et al., 2004) and their hydrophobicity was calculated by the ΔG prediction server v1.0 (https://dgpred.cbr.su.se/index.php?p=home) (Hessa et al., 2007) (for images showed in Figure S1B).

### IMAGING, QUANTIFICATION AND STATISTICAL ANALYSIS

#### Imaging

For detect the subcellular localization of DMA-1 in different mutations, hermaphrodite animals were anesthetized using 5 mM levamisole in M9 buffer (Li et al., 2022; Wang et al., 2021), mounted on 2% agar pads and imaged using Zeiss Cell Observer SD spinning disk confocal microscope equipped with an alpha Plan-Apochromat 63x/1.46 NA objective. Fluorescence signals under excitation light at 488 nm and 561 nm were acquired, respectively (for the image shown in Figures 3A, 5D).

To analyze the secretion of DMA-1SS::moxDendra2::Opsin::FLAG, hermaphrodite animals were anesthetized with 5 mM levamisole /M9 and immobilized on 2% agarose slides for imaging after heat shock treatment at 33 °C for 1 hr and recover at 20 °C for 8 hrs. Fluorescence images were acquired using Zeiss Cell Observer SD spinning disk confocal microscope equipped with alpha Plan-Apochromat 63x/1.46 NA objectives under excitation light at 488 nm and 561 nm, respectively (for the image shown in Figures 1B, 5B,6B, S6A, and S7C).

To determine the subcellular localization of LRP8SS::sfGFP::Opsin::FLAG, two groups of 5 x 10^5^ HEK293FT WT cells or 8 x 10^5^ HEK293FT *ATP13A1* KO cells on Glass Bottom Dish (FEIYUBIO, D35- 20-1-N) were cotransfected according to Figures 2A and 3B. The next day, the medium was changed to PBS with or without 1 μM digitonin and treat for 1 min at room temperature. After incubation, change the medium with 2 mL PBS and image the cells with Zeiss Cell Observer SD spinning disk confocal microscope was equipped with an alpha Plan-Apochromat 63x/1.46 NA objective for fluorescence signals under excitation light at 405 nm, 488 nm, and 561 nm, respectively (for images showed in Figures 2A and 3B).

#### Proteomic Sample Preparation and Data Analysis

For the 4 groups of samples (#1-#4) in Figure S4A, each is about 2 mL cell pellet (from ten 10 cm dishes). #1-#3 were transfected with control, LRP8SS::sfGFP::Opsin::FLAG::His, and PAT- 3SS::sfGFP::Opsin::FLAG::His in HEK293FT cells, respectively, and #4 was transfected with LRP8SS::sfGFP::Opsin::FLAG::His in HEK293FT *ATP13A1* KO cells. Cells were washed twice with cold PBS and collect cell pelleted. Resuspend cells with 8 ml Lysis buffer (50 mM Tris-HCl pH 6.5, 150 mM NaCl, 1 mM EDTA) containing 1% Digitonin and 1X Protease Inhibitor Cocktail and incubate on ice for 1 hr. After incubation, the cells were centrifuged at 10,000 x g for 10 min and the supernatant was transferred to 15 mL centrifuge tube containing ANTI-FLAG M2 Affinity Gel, and then incubate in a cold room for 4-6 hrs. The samples washed with wash buffer (50 mM Tris-HCl pH 6.5, 500 mM NaCl, 1 mM EDTA, 0.03% Digitonin) 4 times (5 mL buffer and 10min/times), and then wash 3 times with PBS. The samples elute by incubating with 3X FLAG Peptide (Sigma-Aldrich) at 4 °C for 1 hr. Eluted samples were sent to the MS platform of the National Center For Protein Science (ShangHai) for MS/MS analysis.

After the mass spectrometry results were relatively quantified by the Label Free Quantitation (LFQ) algorithm, the LFQ intensity of #2-#4 was subtracted from #1 to remove the non-specific result caused by Anti-FLAG beads (it was found that #3 had the similar mass spectrometry results as negative control #1, and no further analysis was performed). Analysis of differentially expressed genes (log_2_FC) for #3 and #4 showed that, compared with *ATP13A1* KO cells, LRP8SS::sfGFP::Opsin::FLAG::His could specifically enrich GET3 in wild-type cells (for images showed in Figure S4A).

#### Quantification and Statistical Analysis

For western blot, ImageJ Software (NIH) was used to analyze the grayscale values from three biological replicates. Student’s t test was used and data are presented as mean ± SEM. Statistical significance is indicated as, ns, not significant; *, p < 0.05; **, p < 0.01; ***, p < 0.001; ****, p < 0.0001.

## ACKNOWLEDGEMENTS

We thank Drs. Yanfen Liu, Yu Cao, Zairong Zhang, and Yingchuan Qi for reagents. Zhaomei Shi and Dr. Piliang Hao from the Multi-Omics Core Facility, Xiaoming Li from the Molecular Imaging Core Facility, and Dr. Yin Xiong from the Molecular and Cell Biology Core Facility in the School of Life Science and Technology at ShanghaiTech University provide technical support. Some strains were provided by the CGC, which is funded by the NIH Program (P40 OD010440). This study was financially supported by the National Natural Science Foundation of China (No. 32170974) and the Natural Science Foundation of Shanghai (No. 22ZR1442400).

## AUTHOR CONTRIBUTIONS

Conceptualization, Y.Z., Z.F., and X.Y.; Investigation, X.Y., Z.F., T.L., and Y.Z.; Data analysis, X.Y., Z.F., T.L., and Y.Z.; Writing, Y.Z. and X.Y.; Review and Editing, Y.Z. and Z.Z.; Funding Acquisition, Y.Z.; Supervision, Y.Z.

## Notes

### Competing Interest Statement

The authors have declared no competing interest.

