## Supplemental Figures for "ATP13A1 engages GET3 to facilitate substrate-specific translocation"

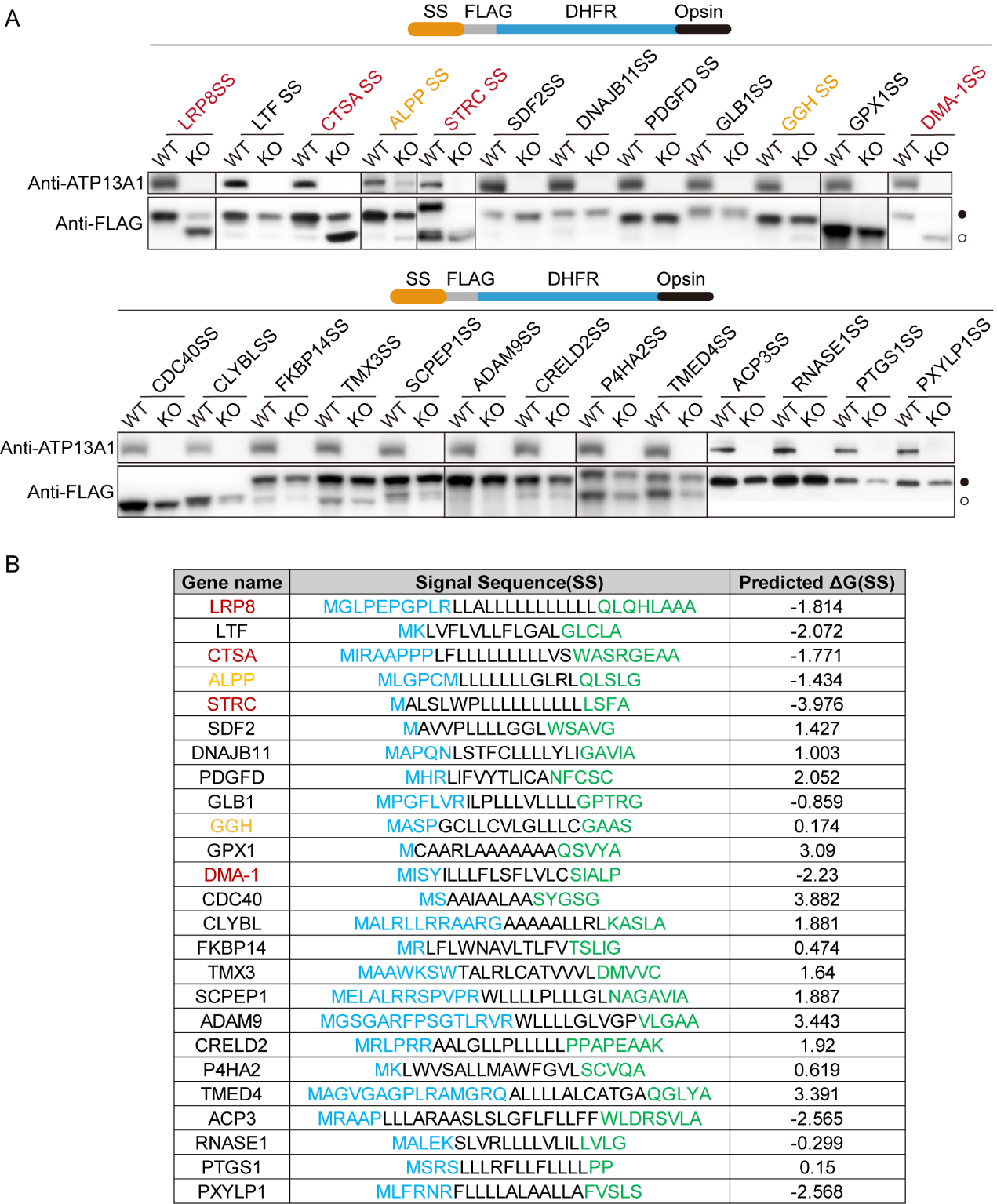

**Figure S1. ATP13A1 is required for SS-mediated ER translocation.**

(A) Translocation efficiency of a variety of SSs assayed in WT or *ATP13A1* KO cells. DHFR is used to generate a protein at a suitable size and to increase protein solubility. Opsin tags containing N-glycosylation sites indicate whether the protein enters into the ER. The SSs whose translocations are highly and moderately dependent on ATP13A1 are shown in red and orange, respectively. Black and clear circles indicate the N-glycosylated and the unglycosylated proteins, respectively.

(B) Sequences and hydrophobicity of SSs assayed in (A). SSs were predicted using SignalP 5.0 (<http://www.cbs.dtu.dk/services/SignalP/>). Their hydrophobicities were calculated by the DG prediction server v1.0 (<https://dgpred.cbr.su.se/index.php?p=TMpred>).

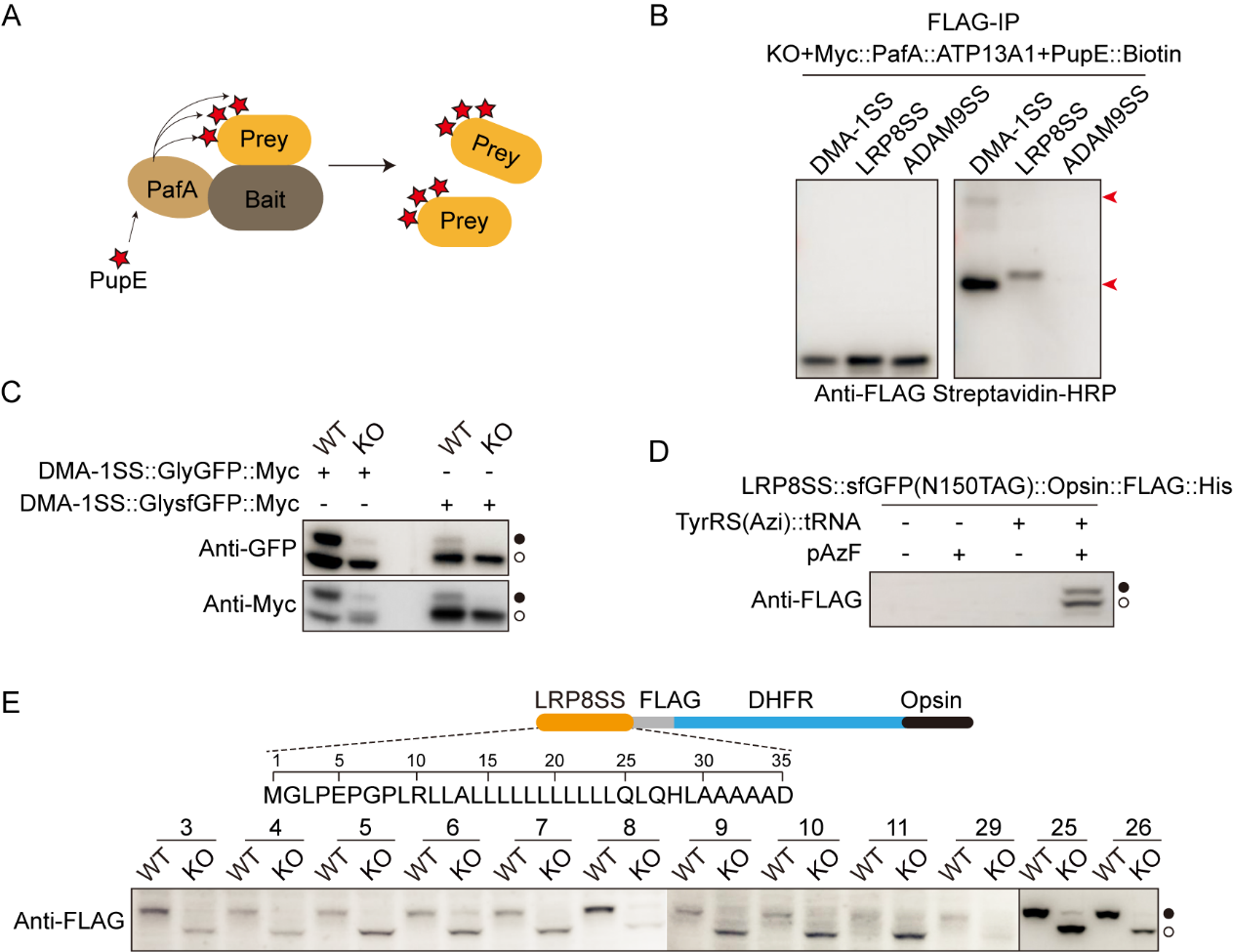

**Figure S2. ATP13A1 directly interacts with SSs.**

(A) Diagram of the PUP-IT proximity tagging system. The bait protein fused with the proximity ligase PafA labels PupE to the prey proteins.

(B) Western blot showing LRP8SS and DMA-1SS, but not ADAM9SS, are labeled with PupE-Biotin by PafA-ATP13A1. The red arrowheads indicate PupE-labeled bands.

(C) sfGFP reduces DMA-1SS translocation. The N-glycosylation site is engineered into GFP (GlyGFP) or sfGFP (GlysfGFP).

(D) LRP8SS::sfGFP(150TAG)::Opsin::FLAG::His is expressed by supplying pAzF and TyRS(Azi)::tRNA.

(E) Screening of optimal positions for incorporating pAzF into LRP8SS. Inserting pAzF into LRP8SS by an amber stop codon, LRP8SS(25TAG), affects neither the expression level nor the P5A-dependence of LRP8SS.

Black and clear circles in (C)-(E) indicate N-glycosylated and unglycosylated proteins, respectively.

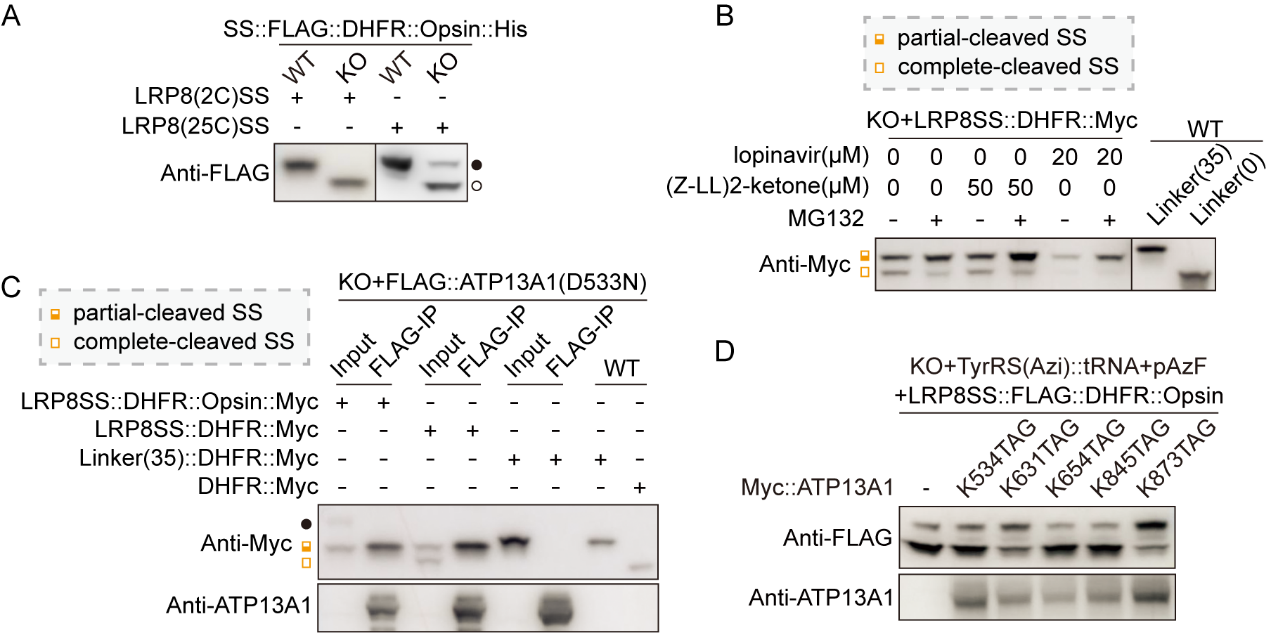

**Figure S3. ATP13A1 interacts with half-cleaved SS and facilitates its translocation.**

(A) LRP8(2C)SS and LRP8(25C)SS with cysteine inserted at position 2 and position 25, respectively, require ATP13A1 for their ER translocation. Black and clear circles indicate N-glycosylated and unglycosylated proteins, respectively.

(B) Partial-cleavage of LRP8SS is not affected in *ATP13A1* KO cells treated with the SPP inhibitor (Z-LL)_2_-ketone or the Ste24 inhibitor lopinavir, given that no larger band showed when incubating with inhibitors.

(C) ATP13A1(D533N) co-immunoprecipitated unglycosylated LRP8SS::DHFR::Opsin::Myc or partial cleaved LRP8SS::DHFR::Myc.

(D) pAzF incorporated into ATP13A1 at position K534, K631, K654, and K845, but not K873, disrupts ATP13A1-mediated LRP8SS translocation.

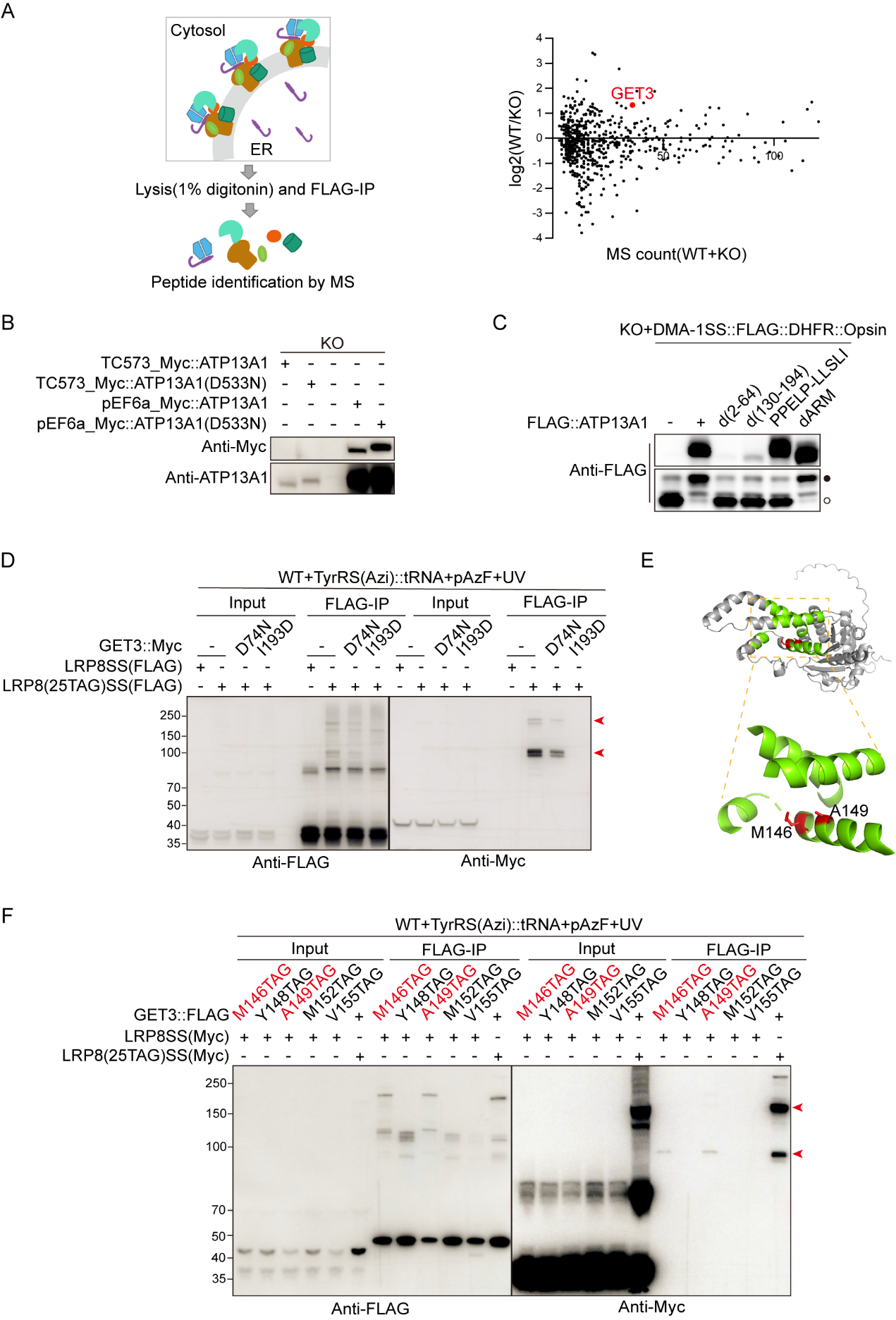

**Figure S4. GET3 is engaged in the ER translocation of N-cleaved SS.**

(A) Diagram showing MS analysis to identify proteins specifically interacting with P5A-dependent SS. Volcano plot showing log_2_ fold change of protein levels in WT compared to *ATP13A1* KO cells pulled down by LRP8SS::sfGFP::Opsin::FLAG::His.

(B) Low- and high-level expression of ATP13A1 variants by the TC573 and pEF6a vectors.

(C) PPELP-LLSLI mutation disrupts ATP13A1-mediated translocation. Black and clear circles indicate the N-glycosylated and the unglycosylated proteins, respectively.

(D) LRP8SS crosslinks with GET3, which is disrupted by GET3(I193D) but not GET3(D74N). LRP8(25TAG)SS::sfGFP::Opsin::FLAG or LRP8SS::sfGFP::Opsin::FLAG was co-transfected with GET3::Myc in WT cells. Red arrows represent the GET3 bands crosslinked with LRP8SS.

(E) Diagram showing the hydrophobic groove in GET3 (highlighted in green).

(F) LRP8SS crosslinks with the residues M146 and A149 in the hydrophobic groove of GET3. Red arrows represent the GET3 bands crosslinked with LRP8SS.

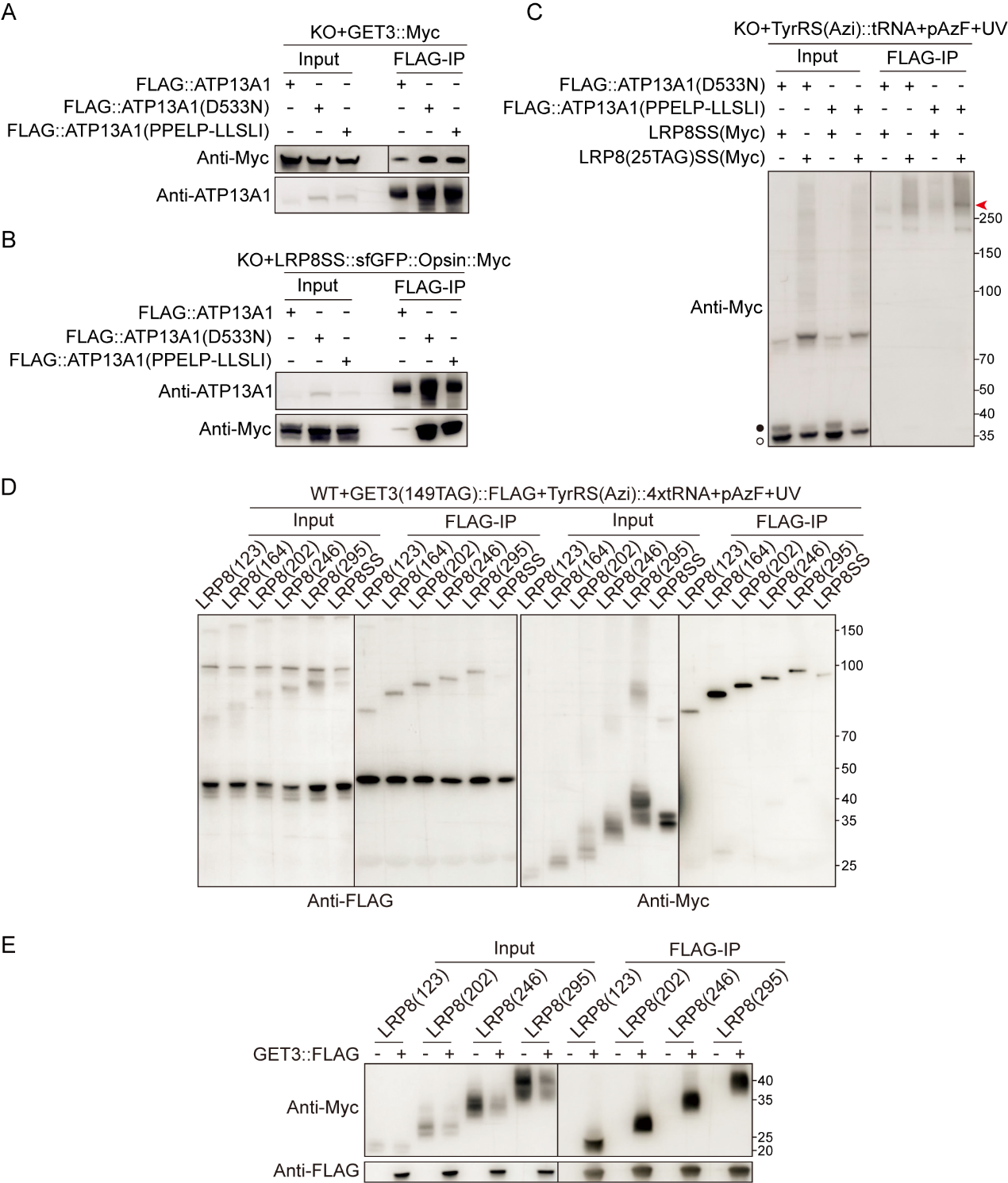

**Figure S5. ATP13A1 engages GET3 for SS translocation.**

(A) ATP13A1 co-immunoprecipitated with GET3, which is not affected by D533N and PPELP-LLSLI mutations in ATP13A1.

(B) ATP13A1 co-immunoprecipitated with LRP8SS, which is enhanced by D533N and PPELP-LLSLI mutations in ATP13A1.

(C) ATP13A1 variants, D533N, and PPELP-LLSLI crosslinks with LRP8SS. Black and clear circles indicate the N-glycosylated and the unglycosylated proteins, respectively. The red arrow represents the band of ATP13A1 variant crosslinked with LRP8SS.

(D) LRP8 truncations at different lengths (123, 164, 202, 246, and 295 amino acids) crosslinked with GET3. GET3(149TAG)::FLAG was co-transfected with different LPR8::Myc variants into WT cells.

(E) Myc-tagged LRP8 truncations at different lengths (123, 164, 202, 246, and 295 amino acids) co-immunoprecipitated with GET3::FLAG. GET3::FLAG was co-transfected with different LPR8::Myc variants into WT cells.

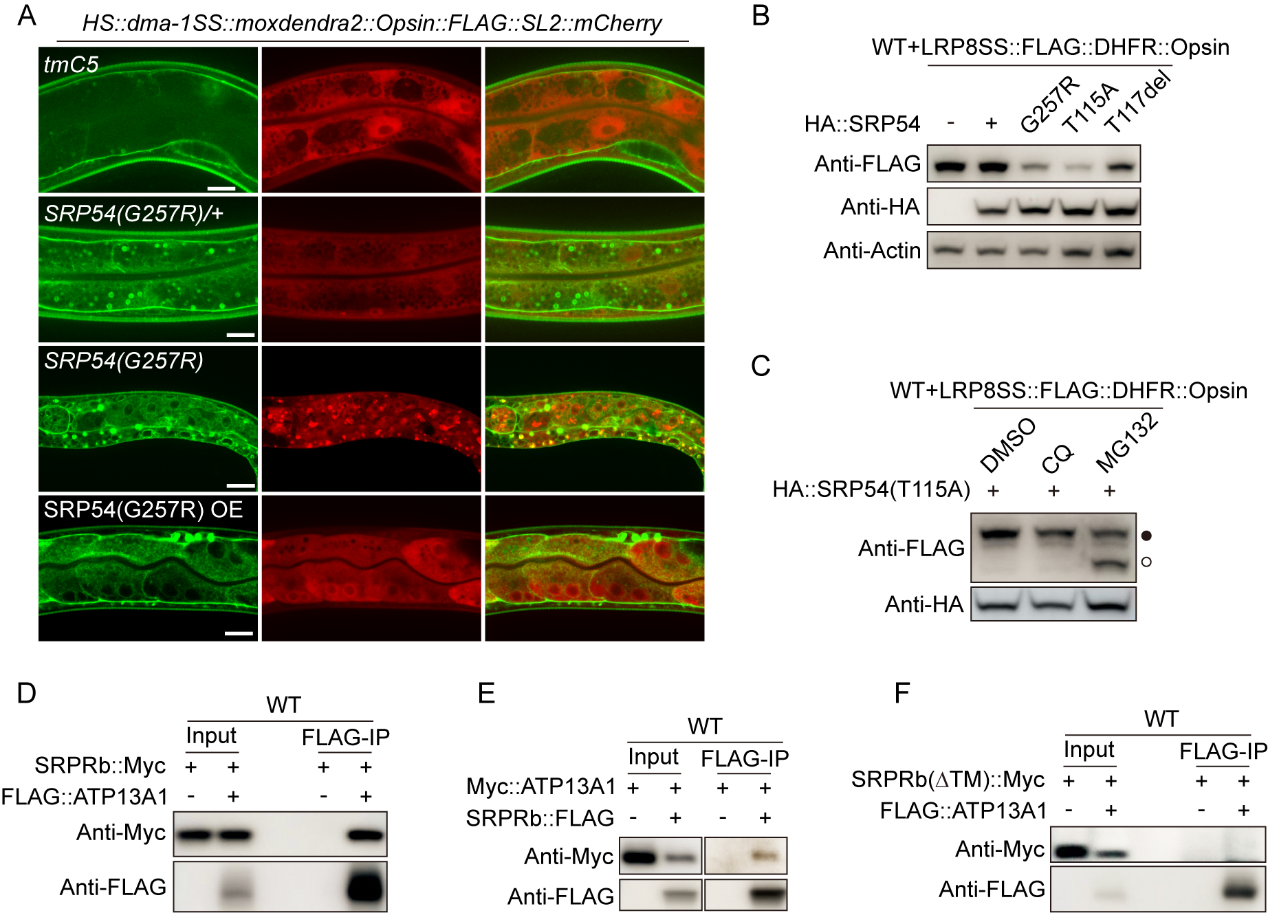

**Figure S6. P5A-dependent SS targets the ER via the SRP complex.**

(A) SRP54(G257R) is a dominant negative mutation which reduces DMA-1SS secretion in *C. elegans*. Scale bars, 10μm.

(B) Overexpression of SRP54(G257R), SRP54(T115A), or SRP54(T117del) reduces LRP8SS level in HEK293FT cells.

(C) SRP54(T115A) overexpression causes LRP8SS degradation, which is stabilized by the protease inhibitor MG132. Black and clear circles indicate the N-glycosylated and the unglycosylated proteins, respectively.

(D) and (E) SRPR co-immunoprecipitated with ATP13A1.

(F) ATP13A1 does not co-immunoprecipitate with SRPR(∆TM).

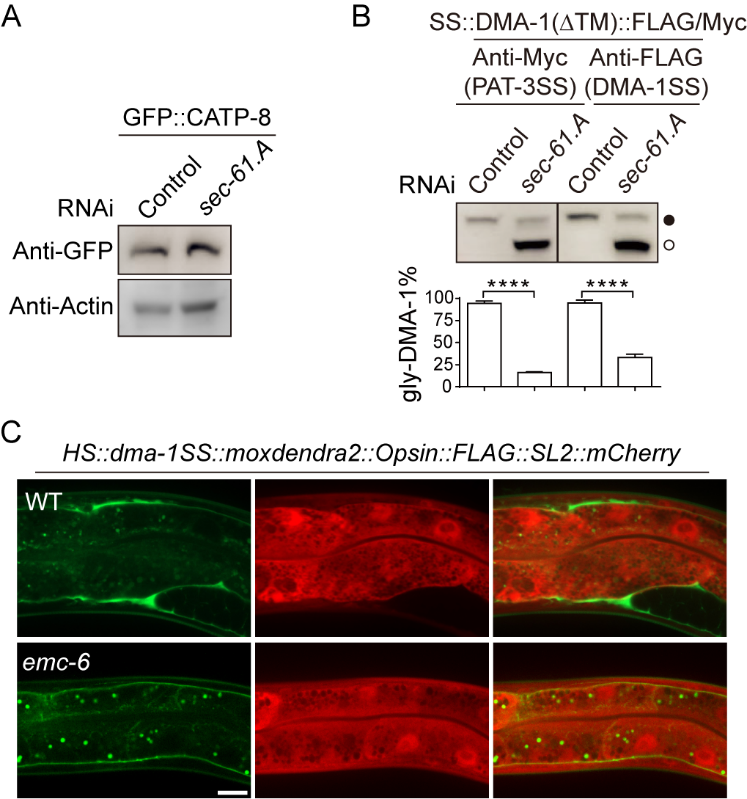

**Figure S7. ER translocation of P5A-dependent SS** **requires SEC61 translocon.**

(A) CATP-8 protein level in GFP::CATP-8 knock-in worms is not affected by *sec-61.A* RNAi.

(B) *sec-61.A* RNAi reduced the ER translocation of both PAT-3SS and DMA-1SS in *C. elegans* at similar levels. Quantifications are from 3 biological replicates and presented as means ± SEMs; *****p*<0.0001 (Student’s *t*-test). Black and clear circles indicate the N-glycosylated and the unglycosylated proteins, respectively.

(C) Secretion of DMA-1SS reporter is not affected in *emc-6* mutants. Scale bar, 10 μm.
